# A conserved disordered region of a splicing factor promotes histone H2B ubiquitination by opposing SUMO-targeted degradation

**DOI:** 10.1101/2025.06.18.660225

**Authors:** Amit Paul, Tracy L. Johnson

## Abstract

While chromatin’s influence on co-transcriptional splicing is well established, the reciprocal model, that spliceosomal components actively shape chromatin, has lacked mechanistic support. Here we show that the C-terminal intrinsically disordered region (IDR) of the yeast splicing factor Prp45 (mammalian SKIP) promotes histone H2B ubiquitination, a hallmark of active transcription. The Prp45 IDR binds and stabilizes Lge1, the scaffold of the H2B ubiquitin ligase complex, enabling its function with the ubiquitin ligase Bre1. When this interaction is disrupted, Lge1 is degraded by the SUMO-targeted ubiquitin ligase Slx5, Bre1 is mislocalized, H2B ubiquitination is lost, and cells become enlarged. Deleting *SLX5* restores Lge1 stability, H2B ubiquitination, and normal cell morphology. Although its sequence diverges across eukaryotes, the IDR’s structural properties are conserved, and the disordered regions of Arabidopsis and human SKIP functionally replace the yeast domain. These findings reveal a conserved IDR-mediated mechanism by which a splicing factor directs chromatin modification.

## Introduction

Since the discovery of introns, studies of the spliceosome—the multi-subunit complex that recognizes and removes these noncoding sequences—have shown that it assembles onto pre-messenger RNA in a stepwise manner. Much of this assembly occurs co-transcriptionally, as the RNA is synthesized by RNA polymerase II on a chromatin template. A critical step in spliceosome assembly is the formation of the activated complex when the Prp19 complex (NTC) and NTC-associated factors (Prp45, Prp46, etc.) bind to the tri-snRNP complex.^1,2,3^ The NTC plays a crucial role in mediating spliceosome rearrangements necessary for catalysis.^2,4^ Both sequence predictions and the high-resolution Cryo-EM structure of the isolated activated complex reveal that one of the NTC-associated factors, Prp45, contains an intrinsically disordered C-terminal region (IDR) that extends out of the complex,^5^ perhaps serving as a site for interaction with other splicing factors during stepwise assembly or other proteins in the proximity of the spliceosome.

Prp45 is essential for cell viability. Nonetheless, the cell can tolerate extensive truncation of Prp45 at its C-terminal end.^6^ A truncation retaining only the first 169 of the 379 residues—hereafter Prp45ΔCTR—supports normal growth at 30°C but is temperature-sensitive, growing poorly at 37°C.^7^ During splicing, this C-terminal region promotes interaction with the second-step helicase Prp22 and thus the fidelity of 3’ splice-site selection ^8^; its loss impairs splicing of a subset of genes.^7^

We previously showed that co-transcriptional assembly of the spliceosome at the step of Prp45 association is influenced by transcription elongation and chromatin state. During transcription elongation, the Set2 histone methyltransferase methylates histone H3 at lysine 36 (H3K36me), a mark of active transcription.^9–11^ H3K36 methylation is recognized by the chromodomain-containing protein Eaf3 ^12–14^, which, in turn, stabilizes the co-transcriptional association of Prp45 and the recruitment of the Prp19 complex. These interactions facilitate co-transcriptional splicing.^13^ As this example illustrates, there is strong evidence that histone modification affects RNA splicing, a relationship that is intuitive given the spatial and temporal proximity of the two reactions. The reciprocal relationship—that splicing factors, in turn, shape chromatin—has also been described, particularly in metazoans.^15,16^ However, direct mechanisms by which splicing factors affect chromatin modification have been difficult to determine. Prp45 is an intriguing candidate for such a role, given its activity at the interface of the two reactions.

The homolog of Prp45, SKIP (Ski-interacting protein), in *Arabidopsis thaliana* and *Homo sapiens* has been shown to play a role in transcription.^17,18^ SKIP interacts with the Paf1 complex and activates FLC transcription to control flowering in *Arabidopsis*.^18,19^ In humans, SKIP associates with the Positive Transcription Elongation Factor b (pTEFb) to facilitate transcription elongation by HIV-1 Tat.^17^ While a role in transcription has not been clearly demonstrated in yeast, the yeast protein Prp45, when fused to the DNA-binding domain, can activate transcription.^6^

H2B monoubiquitination is an important mark that modulates the global chromatin landscape and acts upstream of other key histone modifications associated with transcription elongation, such as H3K4 methylation ^20,21^ and H3K36 methylation.^22,23^ Bre1 is an E3 ligase that mono-ubiquitinates histone H2B at lysine 123 in the coding region during pol II elongation, in collaboration with the E2 conjugating enzyme Rad6.^20,24^ A third protein, Lge1, which was originally identified in a screen for gene deletions that lead to a Large (LarGE) cell phenotype,^25^ is essential for Bre1 catalytic activity. Although Lge1 is largely unstructured, the coiled-coil (CC) domain of Lge1 interacts with the N-terminal region of Bre1, and this interaction is necessary for Bre1-mediated H2BK123 ubiquitination in yeast.^26^ Proper distribution of histone H2B monoubiquitination is necessary for gene activation and is regulated by cycles of deubiquitination. The deubiquitinase Ubp8, which is part of the SAGA complex, deubiquitinates H2B, particularly at the 5’ end of genes.^22^ While Ubp8 is a well-characterized downstream regulator of H2B ubiquitination, no direct regulator of the E3 ligase complex itself is known. Since the activity of the H2B ubiquitination complex is dependent on interactions between the components Bre1 and Lge1, this interaction is a potential target of regulation—raising the possibility that the mark is controlled by a factor acting from outside the ligase complex.

Here, combining genetics, proteomics, and biochemistry, we show that the C-terminal IDR of Prp45 binds and stabilizes Lge1, the scaffold of the H2B ubiquitin ligase complex, enabling H2B monoubiquitination together with the ubiquitin ligase Bre1. When the IDR is removed, Lge1 is degraded in a manner dependent on the SUMO-targeted ubiquitin ligase (STUbL) Slx5, H2B ubiquitination is reduced, and cells become enlarged; deleting SLX5 restores Lge1 stability, H2B ubiquitination, and normal cell morphology. These findings connect the two questions raised above—how a splicing factor directs a chromatin mark, and how the H2B ligase complex is itself controlled—by placing the Prp45–Lge1 interaction at the center of ubiquitin–SUMO crosstalk.

Although its sequence diverges across eukaryotes, the structural properties of the C-terminal IDR of Prp45/SKIP are conserved, including approximately 50 amino acid regions with propensity to form helical structure. In fact, the IDR of *Arabidopsis* and human SKIP functionally replace the yeast domain and recapitulate the chromatin modification and cell morphology phenotypes, demonstrating the functional conservation of the C-terminal IDR in Prp45/SKIP. These studies reveal how an RNA splicing factor through its IDR prevents SUMO-targeted degradation to promote H2B ubiquitination.

## Results

### The C-terminal portion of Prp45 is required for optimal cell viability

Previous studies provided evidence that Prp45 is recruited co-transcriptionally and in close proximity to the chromatin.^13^ Prp45 contains an SNW domain (amino acids 1-169) that is essential for viability. The C-terminal region encompasses a helical domain and the SH2-like domain, which, in isolation, is predicted to be highly disordered, as illustrated by analysis using AIUPred ^27^ (Fig. 1a).

**Figure 1.**
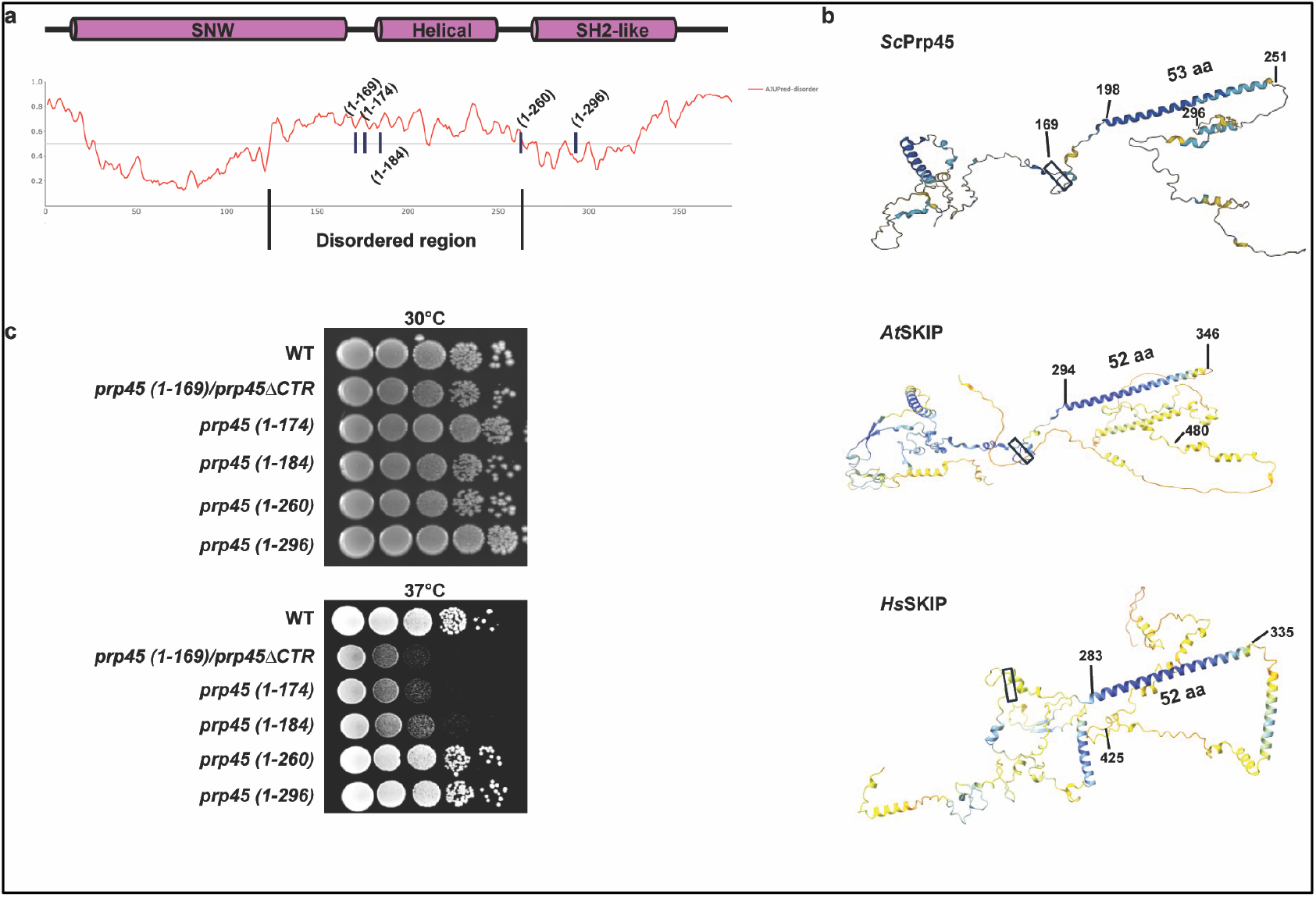
Prp45 has a disordered C-terminus that is required for viability at elevated temperatures. (a) Schematic of the domain architecture of the Prp45 protein. The disordered region of the protein was predicted by AIUPred. (b) The structure of *Sc* PRP45, *At* SKIP, and *Hs* SNW1 protein generated by AlphaFold showing the conserved alpha helical domain. (c) The C-terminal mutants of Prp45 shows a temperature dependent growth defect. Prp45 C-terminal truncated mutants were grown at 30°C and serially diluted on YPD and incubated at 30°C or 37°C (four days each).

The homologs of Prp45 in *Arabidopsis* (*At* SKIP) and Humans (*Hs* SKIP) are also spliceosome components. Although the amino acid sequence of the C-terminus is different from yeast (Fig. S1), they share a striking structural similarity (Fig. 1b). Interestingly, a region of the C-terminus with conserved alpha-helical propensity is thought to be a site of protein-protein interactions and is approximately 52 amino acids long across species (Fig. 1b).

To begin to evaluate the function of this disordered, C-terminal region of the protein, we generated truncation mutants of Prp45; *prp45 (1-169)*, *(1-174)*, *(1-184)*, *(1-260)* and *(1-296)*, where the residues after 169, 174, 184, 260, and 296 were deleted, and the cells were grown at 30°C and 37°C (Fig. 1c). The largest truncation, which eliminates the helical domain and a linker region between the SNW and the helical domain, *prp45 (1-169)* or *prp45ΔCTR*, shows the most dramatic growth defect at elevated temperature. The strains with the second-largest growth defects, *prp45 (1-174)* and *prp45 (1-184)*, eliminate the helical region while retaining the linker region. *prp45 (1-260)* and *prp45 (1-296)*, containing the region beyond the predicted disordered regions, grow similarly to wild-type cells at 37°C (Fig. 1c). These results suggest that the disordered region of Prp45 that has a propensity to form helical structure is important for its function, warranting a more detailed exploration of the domain’s protein-binding properties and potential interactors.

### The Prp45 C-terminus interacts with the H2B ubiquitination machinery and is critical for H2B ubiquitination

To identify specific proteins that interact with Prp45 via the C-terminal region, proximity labeling was performed. Briefly, biotin ligase BirA was added to the C-terminal region of either full-length Prp45 or Prp45ΔCTR, and the strains were grown in the presence of biotin. The biotinylated proteins were immunoprecipitated using streptavidin beads, followed by mass spectrometry (Fig. S2a). Full-length Prp45 interacts with 251 different proteins. Of these, 213 proteins show differential interactions when the C-terminal end is truncated (Fig. S2b). We first focused on those with a significant decrease in interactions, then on those in which interactions were lost upon truncation of the C-terminal end. Not surprisingly, most of the abrogated interactions are with known splicing factors, such as Prp19, a fellow member of the NTC. The interactions between Prp45 and Brr2, Clf1, Prp46, and Spp2 are significantly reduced in Prp45ΔCTR, compared to the wild type (Fig. S2c).

Thirty-eight proteins are not detected at all when the C-terminus is truncated, suggesting that their interactions with Prp45 are dependent on the C-terminus (Table S1). Hierarchical clustering of the 38 proteins reveals that the largest class of proteins is involved in RNA splicing. Nevertheless, an intriguing subset is implicated in transcription elongation and chromatin function (Table S1). One of the interactions lost when the C-terminus of Prp45 is deleted is with Bre1, the E3 ligase that, together with its E2, Rad6, monoubiquitinates histone H2B.^28,29^ Monoubiquitination of H2B requires a third component of this complex, Lge1, which interacts directly with Bre1 and is required for Bre1’s H2B-ub activity.^30,31^

To determine if there is a functional relationship between H2B monoubiquitination and the Prp45 C-terminus, *BRE1*, *RAD6*, and *LGE1* were each deleted in combination with *prp45ΔCTR,* and the strains were grown at 30°C and 37°C. While growth of the double mutants is not significantly impaired at 30°C, at 37°C, *prp45ΔCTR bre1Δ*, *prp45ΔCTR rad6Δ*, and *prp45ΔCTR lge1Δ* all show a growth defect compared to *prp45ΔCTR*; for *prp45ΔCTR bre1Δ* and *prp45ΔCTR lge1Δ*, growth is barely visible (Fig. 2a).

**Figure 2.**
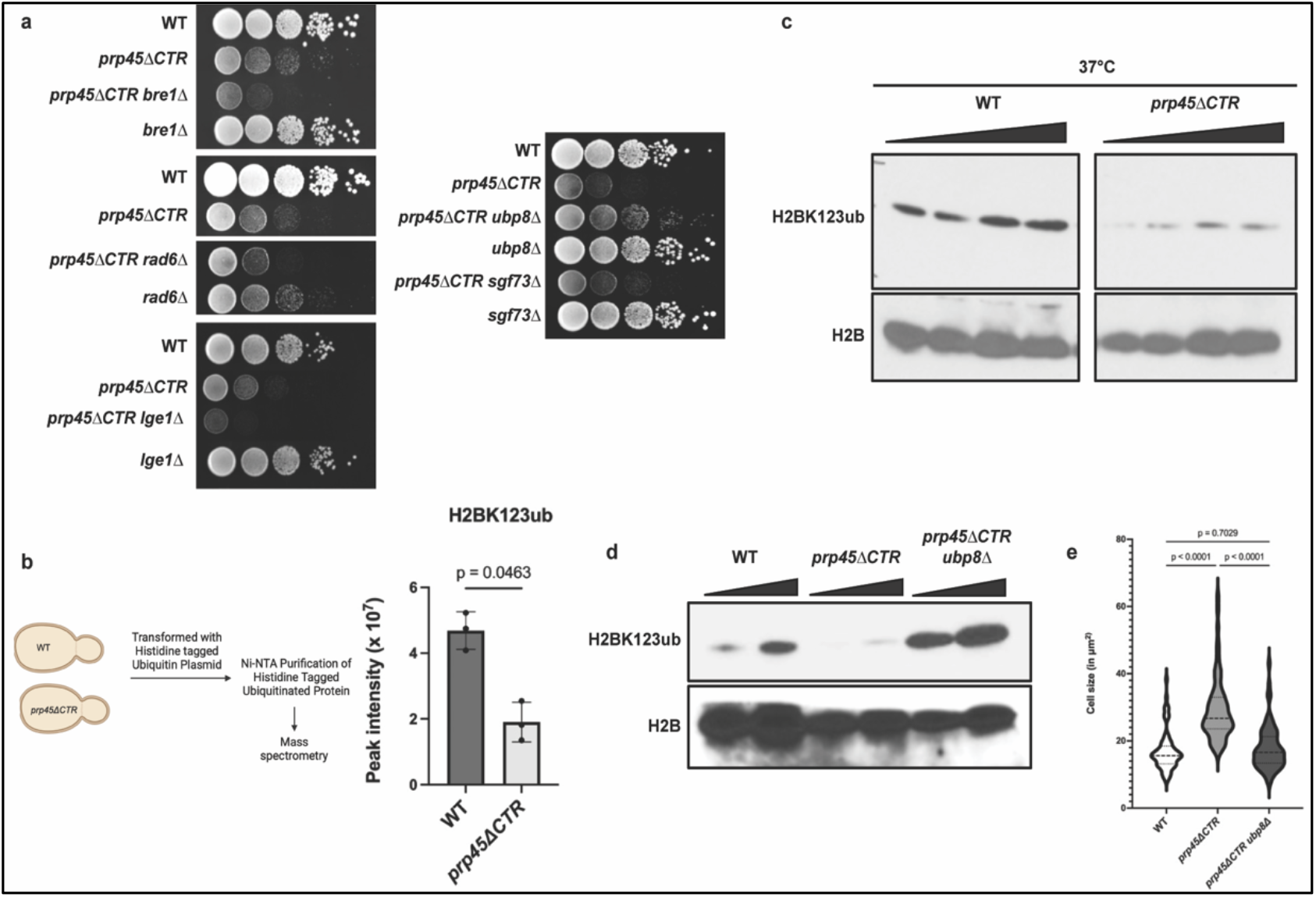
The Prp45 C-terminus interacts with the histone ubiquitination machinery. (a) H2B-ub ligase complex deletions (*bre1Δ*, *rad6Δ*, *lge1Δ*) exacerbate the *prp45ΔCTR* growth defect; deletion of H2B deubiquitinase complex components (*ubp8Δ* and *sgf73Δ*) suppresses it. Cells were serially diluted tenfold, plated on YPD, and grown at 37°C for four days. (b) Schematic of *in vivo* ubiquitome analysis using WT or *prp45ΔCTR* cells. Quantification of ubiquitinated H2BK123 peptides in WT and the *prp45ΔCTR* strain reflects the results from the mass spectrometry data. Model was generated using BioRender. (c) Truncation of the C-terminal domain of Prp45 causes a significant reduction in H2B monoubiquitination. Immunoblot analysis of ubiquitinated histone H2B in whole-cell extracts isolated from WT and *prp45ΔCTR* strains grown at 37°C. Increasing amounts of lysates were loaded onto 15% SDS-PAGE, and ubiquitinated H2B, or H2B, was detected with anti-H2BK123ub or anti-H2B antibodies. (d) Deletion of *UBP8* restores the H2BK123 ubiquitination. H2BK123 monoubiquitination was detected from WCE of WT, *prp45ΔCTR,* and *prp45ΔCTR ubp8*Δ strains. (e) The large cell size of the *prp45ΔCTR* strain is reversed by the deletion of *UBP8*. Confocal imaging (DIC) of WT, *prp45ΔCTR,* and *prp45ΔCTR ubp8Δ* strains grown to OD_600_ 0.6 at 37°C was performed, and cell size measured using ImageJ. Cell size (n = 80) was plotted in GraphPad PRISM, and an unpaired one-way ANOVA with Tukey’s post hoc test was used to calculate *p*-values.

Since mutations that abrogate H2B monoubiquitination show strong negative genetic interactions with Prp45, we predicted that mutations that stabilize monoubiquitination would reverse the *prp45ΔCTR* growth defect. Ubp8 and Sgf73 are members of the SAGA complex and contribute to histone H2B deubiquitination.^32–35^ Deletion of either *UBP8* or *SGF73* in the *prp45ΔCTR* strain partially rescues the growth defect at 37°C (Fig. 2a, right image), with deletion of *UBP8*, the gene encoding the catalytic protein, showing the strongest suppression.

Considering the functional and physical interactions between H2B ubiquitination machinery and Prp45, H2B ubiquitination was assessed in cells harboring a Prp45 C-terminal truncation. Analysis of the ubiquitinome (Fig. 2b schematic) from wild-type and *prp45ΔCTR* cells reveal a significant reduction in peak intensity of H2BK123 peptides in *prp45ΔCTR* (Fig. 2b). Immunoblot analysis shows a significant reduction in H2BK123 ubiquitination in *prp45ΔCTR* at 37°C. In fact, there is an approximately 90% decrease in H2BK123ub in the *prp45ΔCTR* strain, although the levels of H2B remain unchanged (Fig. 2c). Hence, the C-terminal domain of Prp45 affects H2BK123 ubiquitination, likely by modulating the catalytic activity of Bre1, the ubiquitin ligase.

Since deletion of *UBP8* partially rescues the growth defect in *prp45ΔCTR,* it seemed likely that it would also rescue the H2BK123 ubiquitination defect. Indeed, restoration of H2BK123 ubiquitination is observed in *prp45ΔCTR ubp8Δ* (Fig. 2d), consistent with the observation of a partial rescue of the *prp45ΔCTR* growth defect when *UBP8* is deleted. Notably, although H2B ubiquitination is restored, growth is partially restored, which points to a contribution of the other functions of the C-terminal domain in addition to its role in H2B ubiquitination for optimal growth.

H2B ubiquitination has been shown to be involved in maintaining cell size. Mutations in any of the genes that regulate H2B ubiquitination lead to a LarGE cell phenotype. ^36^ Interestingly, previous observations reveal that when the C-terminus of Prp45 is truncated, cells are also large, elongated, and highly heterogeneous.^8^ In a comparison between the cell sizes of *prp45ΔCTR* and *lge1Δ*, relative to WT cells, *prp45ΔCTR* cells are almost 2.5 times larger than the wild type cells and similar in size to *lge1Δ* cells (Fig. S3 and Fig. 2e). If the aberrant cell size of *prp45ΔCTR* is due to effects on H2B ubiquitination, we predicted that deleting *UBP8* would reverse the cell size defect. Indeed, stabilizing H2B ubiquitination by deleting *UBP8* reverses the elongated-cell phenotype and restores cell size in *prp45ΔCTR* (Fig. 2e).

### The C-terminal domain of Prp45 interacts with and stabilizes the ubiquitin ligase scaffold protein, Lge1

Although Bre1 is the ubiquitin ligase responsible for H2B monoubiquitination, Bre1’s activity is dependent on the Lge1 protein.^37^ A recent analysis of the Lge1-Bre1 interaction reveals that Lge1 phase separates and forms condensates, which is mediated by its intrinsically disordered domain. The coiled-coil (CC) domain of Lge1 interacts with the middle, or Lge1 binding domain (LBD), of Bre1, and this interaction is necessary for Bre1-mediated H2BK123 ubiquitination.^26^ Although Lge1-deleted cells can normally grow at 30°C and 37°C, they are morphologically larger than wild-type cells. The Bre1-Lge1 interaction and function appear to be conserved. A recent study of the mammalian proteins responsible for H2B ubiquitination shows that RNF20/RNF40, the mammalian homolog of Bre1, interacts with the conserved coiled-coil domain of WAC protein (WAC-CT) (Lge1 homolog in mammals), and this interaction is crucial for H2B ubiquitination.^38^

While proximity labeling provides insights into the physical distance between two proteins, it is difficult to assess whether these interactions are direct or indirect since any protein residing within a 10 nm diameter can be biotinylated.^39^ In light of the observation of a Prp45 C-terminus-dependent interaction with Bre1, we sought to capture interactions between these proteins by co-immunoprecipitation, but did not detect a physical interaction between Bre1 and Prp45, suggesting that the interaction observed by proximity labeling may be weak and/or indirect. We therefore considered proteins that are part of the H2B ubiquitination complex to determine whether they interact with Prp45, starting with Lge1, the Bre1 cofactor required for its catalytic activity.^31^ Full-length Prp45 immunoprecipitates Lge1, and the interaction is reduced when the C-terminus of Prp45 is truncated (Fig. 3a).

**Figure 3.**
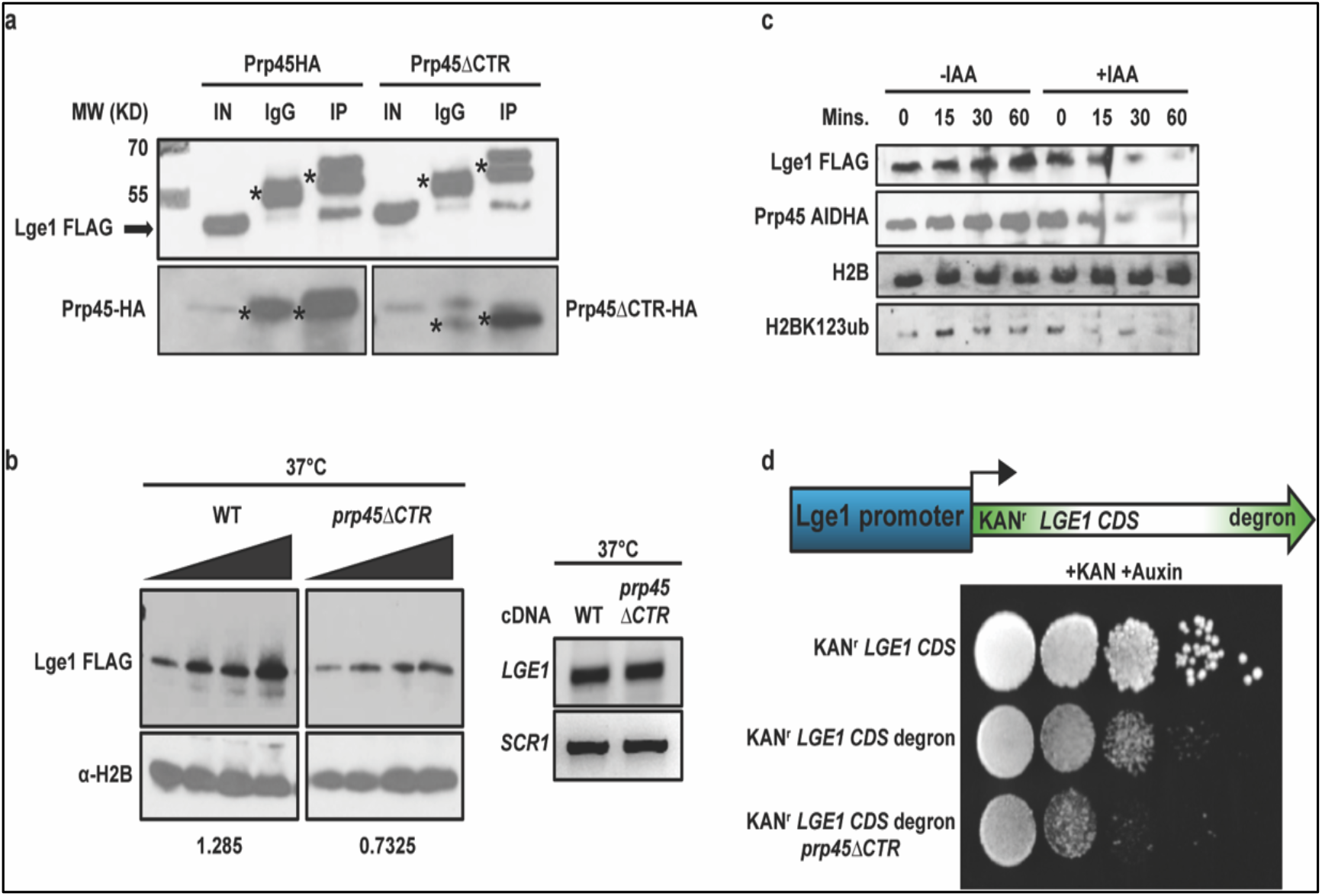
The interaction between Prp45 and Lge1 is required for the stability of the Lge1. (a) Prp45 physically interacts with the Lge1. Prp45-HA was immunoprecipitated with Anti-HA antibody and immunoblotted with Anti-FLAG antibody to detect Lge1. 2% of the total lysate was used as input. Asterisks in experimental lanes 2, 3, 5, and 6 indicate the heavy chains of the primary antibody (top panel) and the light chains (bottom panel left and right). (b) Loss of the C-terminal domain of Prp45 causes a reduction in Lge1 stability. Immunoblot analysis of whole-cell extracts isolated from WT and *prp45ΔCTR* cells grown at 37°C. Lge1-FLAG was detected using an anti-FLAG antibody. Densitometric quantification was performed using ImageJ. RT-PCR analysis of LGE1 RNA at 37°C shows comparable levels of RNA between the wild type and the *prp45ΔCTR* strain (right image). (c) The loss of Prp45 protein leads to Lge1 protein instability. Immunoblot analysis shows Lge1 protein levels upon auxin-inducible degradation of Prp45 when cells were incubated with IAA at different time points (0, 15’, 30’, and 60’). The DMSO control is shown in lanes 1-4. Prp45-AID-HA and Lge1-FLAG was detected using anti-HA and anti-FLAG antibodies, respectively. Ubiquitinated H2BK123 and H2B were detected using anti-H2BK123-ub and anti-H2B antibodies. (d) Growth assay illustrating increased sensitivity to degradation in Lge1 instability when Prp45 is truncated. The entire Lge1 CDS was N-terminally tagged with a Kanamycin resistance cassette and C-terminally tagged with a degron sequence under the native Lge1 promoter in wild-type and *prp45ΔCTR* strains (schematic). Cells were tenfold serially diluted on a YPD + kanamycin (200 μg/ml) + IAA (1 mM) plate and incubated at 30°C.

Lge1 is a highly disordered protein, with most of its residues (~85%) residing in disordered regions, as illustrated in Figure S4a (upper image); it also contains a short, structured coiled-coil (CC) domain through which it interacts with other proteins, such as Bre1.^26^ These data suggest that Prp45’s effects on the catalytic activity of Bre1 are likely through its interactions with Lge1. While we cannot rule out the possibility that Prp45 interacts directly with *both* Lge1 and Bre1 (perhaps weakly in the case of the latter), Prp45 interacts with Lge1, and the Lge1-Bre1 activity is dramatically affected by the Prp45 C-terminus. Based on the known structural information for Prp45 and Lge1, AlphaFold-multimer in Google Colab Fold ^40^ was used to predict protein-protein interacting regions between the proteins. The *in-silico* analysis suggests interactions at the C-terminal region of Prp45 and a structured coiled-coil (CC) domain of the Lge1 protein (Fig. S4a lower image).

In light of reports that Lge1 is a disordered protein, and intrinsically disordered segments can affect protein half-life ^41^, we considered the possibility that Lge1’s interaction with the C-terminus of Prp45 is an important contributor to its stability. To determine if this is the case, immunoblot analysis of Lge1 protein was performed in wild-type and *prp45ΔCTR* strains. Although *LGE1* RNA levels remain unchanged (Fig. 3b, right), the Lge1 protein becomes unstable in the *prp45ΔCTR* strain when compared to the Lge1 protein isolated from wild-type cells, particularly at 37°C (Fig. 3b, left).

To further assess whether Prp45 contributes to Lge1 stability, Lge1 protein levels were analyzed in an auxin-inducible degron strain, where Prp45 was degraded in the presence of IAA over time (0, 15’, 30’, 60’). Indeed, Lge1 levels decrease in a time-dependent manner, with kinetics similar to Prp45 (Fig. 3c). H2BK123 ubiquitination levels are also gradually reduced during the degradation of Prp45 and Lge1, with comparable kinetics. When cells are exposed to DMSO at the same time points, Lge1 protein levels remain unchanged, indicating IAA (and hence Prp45) dependent loss of Lge1 (Fig. 3c), which abrogates H2B ubiquitination.

To demonstrate that the presence of the C-terminal region of Prp45, specifically, protects Lge1 from degradation, we developed a different Lge1 stability assay with a clear phenotypic readout. The entire *LGE1* CDS was tagged with a degron at its C-terminal end, and a Kanamycin resistance marker was included at its N-terminal end under the native *LGE1* promoter. The rate of degradation of Lge1 protein was examined in the wild type or the *prp45ΔCTR* strain by assaying Kan^r^ (Fig. 3d, schematic). When the cells were grown and serially diluted on a YPD + Kanamycin + IAA plate (Fig. 3d), the results show a clear growth difference between the Lge1 degron in the wild-type background vs. the *prp45ΔCTR* strain. Degradation of Lge1 in the presence of IAA causes loss of kanamycin resistance; the *prp45ΔCTR* exacerbates the growth defect, consistent with enhanced Lge1 destabilization when the CTR of Prp45 is absent (Fig. 3d).

These data suggest that the IDR of Prp45 protects Lge1 from degradation. Intriguingly, deletion of the Prp45 IDR leads to lower levels of Lge1 and increased ubiquitination; the decrease in protein relative to the increase in ubiquitination is particularly striking (Fig. S4b) and suggests rapid proteasome-mediated degradation of the Lge1 protein. This led us to identify the ubiquitin ligase complex that targets Lge1.

### The SUMO-targeted ubiquitin ligase Slx5 mediates Lge1 destabilization in the absence of the Prp45 C-terminus

Having established that the Prp45 C-terminus is required for Lge1 stability, we asked what drives Lge1 turnover when the C-terminus is absent. A large-scale Synthetic Genetic Array revealed that the growth defect of the Prp45 allele *prp45-5002*, was suppressed by deletion of the gene encoding Slx5. ^42^ Slx5, along with Slx8, forms the SUMO-targeted ubiquitin ligase (STUbL) that directs SUMO-modified nuclear proteins to proteasomal degradation. ^43^ ^44^ Because Lge1 is a highly disordered protein whose stability depends on its protein interactions, we considered the possibility that Slx5 targets Lge1 for degradation when it is no longer protected by the Prp45 C-terminus. To test this genetically, *SLX5* was deleted in the *prp45ΔCTR* strain. Consistent with the SGA analysis, deletion of *SLX5* suppresses the *prp45ΔCTR* growth defect at 37°C (Fig. 4a). Since Lge1 is associated with chromatin, and the Prp45 C-terminal region protects it from STUbL-mediated degradation, we anticipated that loss of the Prp45 CTR would increase Slx5 at the chromatin. So, we performed a protein transit assay followed by mass spectrometry. The histone protein H2A.Z (encoded by *HTZ1*) was tagged with the biotin ligase in either wild type or the *prp45**Δ**CTR* strain (Fig. 4b, schematic) to look for changes in Slx5 association with chromatin (H2A.Z).

**Figure 4.**
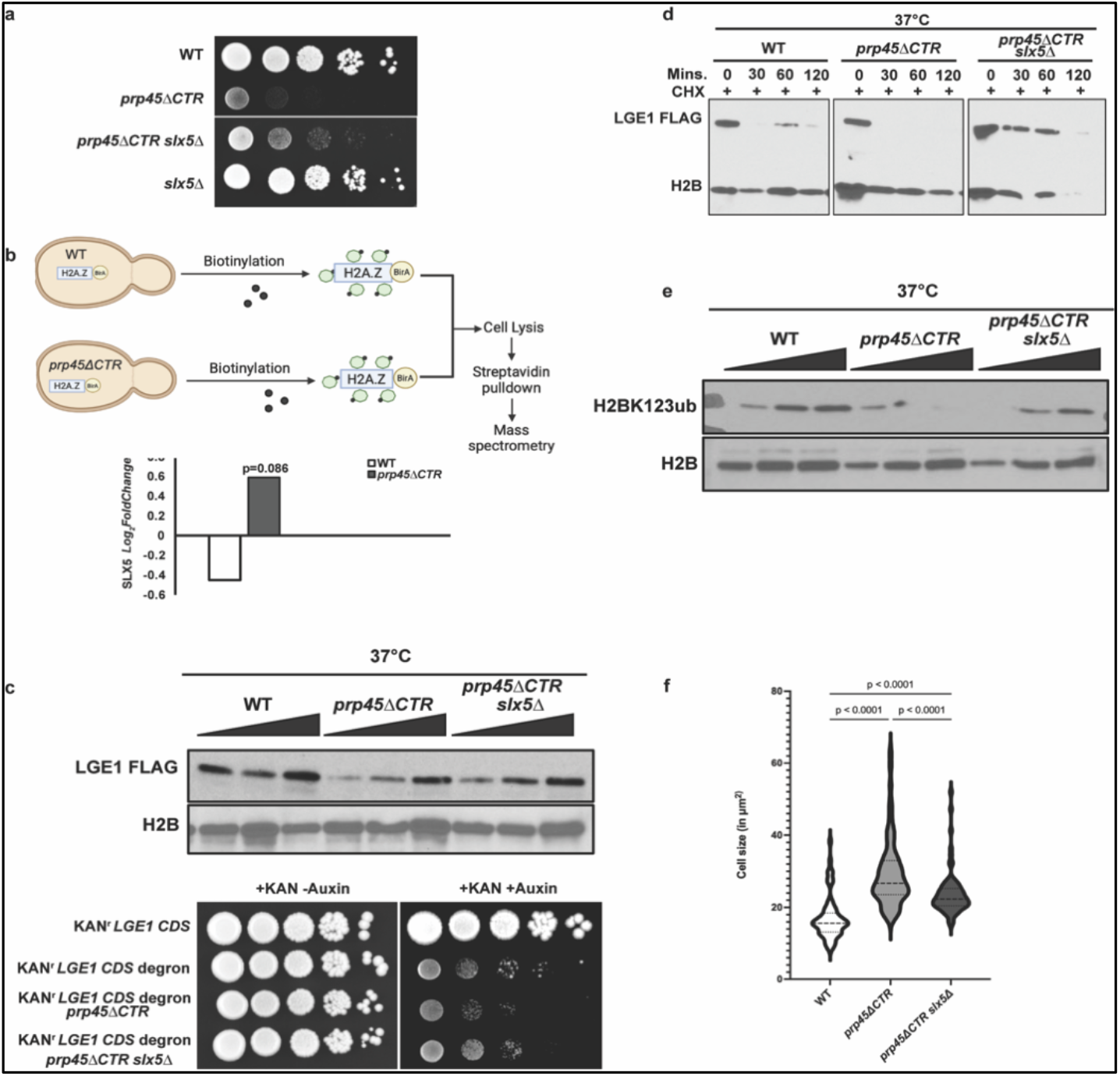
STUbL regulates Lge1 turnover. (a) STUbL E3 ligase deletion rescues the growth defect in 37°C. The strains indicated were serially diluted and spotted on YPD. The plate was kept at 37°C for 3 days before imaging. (b) Slx5 enrichment assay. Modified protein transit assay was performed using H2A.Z as a chromatin bait protein, The enrichment of Slx5 was presented as log2 fold change of wild type vs *prp45ΔCTR.* The adjusted p value was used to show the trend of enrichment. Model was generated using BioRender. (c) Deletion of *SLX5* rescues Lge1 stability. The immunoblot shows whole-cell extracts isolated from WT, *prp45ΔCTR and prp45ΔCTR slx5Δ* cells grown at 37°C. Lge1-FLAG was detected using an anti-FLAG antibody. Growth assay (lower image) illustrates an increase in Lge1 instability when *SLX5* is deleted. Lge1 is N-terminally tagged with a Kanamycin resistance cassette and C-terminally tagged with a degron sequence under the native Lge1 promoter in wild-type, *prp45ΔCTR* and *prp45ΔCTR slx5Δ* strains. Cells were tenfold serially diluted on a YPD + kanamycin + IAA plate and incubated at 30°C. (d) CHX experiment showing the degradation kinetics of lge1 protein. Cells were grown to reach OD-0.6 followed by 100ug/mL Cycloheximide treatment at 37°C. Cells were collected every 30 minutes up to 2 hours. Whole-cell extracts were isolated from WT, *prp45ΔCTR and prp45ΔCTR slx5Δ,* and lge1 was detected using anti-FLAG antibody. (e) Deletion of slx5 restores the H2B ubiquitination. Increasing amounts of lysates were loaded onto 15% SDS-PAGE, and ubiquitinated H2B, or H2B, was detected with anti-H2BK123ub or anti-H2B antibodies in WT, *prp45ΔCTR,* and *prp45ΔCTR slx5*Δ strains. (f) *SLX5* deletion partially rescues cell size. The large cell size of the *prp45ΔCTR* strain is partially rescued by the deletion of Slx5. WT, *prp45ΔCTR,* and *prp45ΔCTR slx5Δ* strains were grown to OD_600_ 0.6 at 37°C and Confocal imaging (DIC) was performed. Cell size was measured using ImageJ. Cell size (n = 80) was plotted in GraphPad PRISM, and an unpaired one-way ANOVA with Tukey’s post hoc test was used to calculate *p*-values.

First, Prp45 is significantly enriched relative to the minus biotin control in the HTZ1-TurboID proximity proteomics, both in wild type and *prp45**Δ**CTR* strains (Log_2_FC = 0.285 and 0.318, respectively), confirming that Prp45 is chromatin-associated and detected by the proximity labeling system. Importantly, Slx5 is, indeed, enriched in the chromatin fraction in the *prp45**Δ**CTR* strain compared to wild type cells (Fig. 4b). We predicted that the increased enrichment of the Slx5 protein at the chromatin in the *prp45**Δ**CTR* strain contributes to the increase in degradation of Lge1. Conversely, deletion of *SLX5* would be predicted to stabilize the Lge1 protein. To test this, Lge1 protein was measured by Western blot, and Lge1 levels were found to be restored in the *prp45ΔCTR slx5Δ* strain compared to *prp45ΔCTR* (Fig. 4c). This was corroborated by the orthogonal degron assay: deletion of *SLX5* in the Kan^r^ Lge1 CDS degron *prp45ΔCTR* strain partially restores growth on IAA + Kan plate (Fig. 4c lower image).

The protective role of the C-terminus of Prp45 was further evaluated by treating cells with cycloheximide (CHX) to stop new protein synthesis and measure Lge1 degradation at 37°C. Lge1 is stable in the absence of the Slx5 protein (Fig. 4d), while *LGE1* RNA levels are unchanged. Restoration of Lge1 is accompanied by recovery of H2BK123 ubiquitination (Fig. 4e) and reversal of the enlarged, elongated-cell morphology characteristic of *prp45ΔCTR* (Fig. 4f).

Together, these data indicate that, in the absence of the Prp45 C-terminus, Lge1 is destabilized through an Slx5-dependent degradation pathway, and that the Prp45 C-terminus protects Lge1 from this turnover. Whether Slx5 targets Lge1 directly—for example, following SUMO modification of Lge1—or acts indirectly on a Lge1-associated factor remains to be determined. Nonetheless, these results demonstrate a novel function of the STUbL complex and a protective function for the IDR of Prp45.

### Loss of Lge1 leads to a decrease in Lge1-dependent nuclear puncta and altered Bre1 distribution

Lge1 was first identified in a screen for genes whose deletion disrupts cell size homeostasis. Although *LGE1* is not essential for cell viability, morphologically, the cells appear to have this large phenotype when *LGE1* is deleted.^25^ Lge1 forms puncta in the nucleus to locally concentrate Bre1.^26^ Since Prp45 degradation leads to unstable Lge1 protein (Fig. 3c), we predicted that puncta formed by the Lge1 protein would also be reduced. To test this prediction, Lge1 was GFP-tagged in the Prp45 degron strain, and cells were grown in the presence of IAA or DMSO, followed by confocal microscopy. Indeed, GFP intensity was significantly reduced in the presence of IAA (Fig. 5a) compared with the DMSO control. Lower GFP intensity was quantified and confirmed using cell pose ^45^ (Fig. 5b right). The quantification also revealed that cells become larger when Prp45 is degraded (Fig. 5b left), similar to the phenotype observed when H2B ubiquitination is reduced.^36^

**Figure 5.**
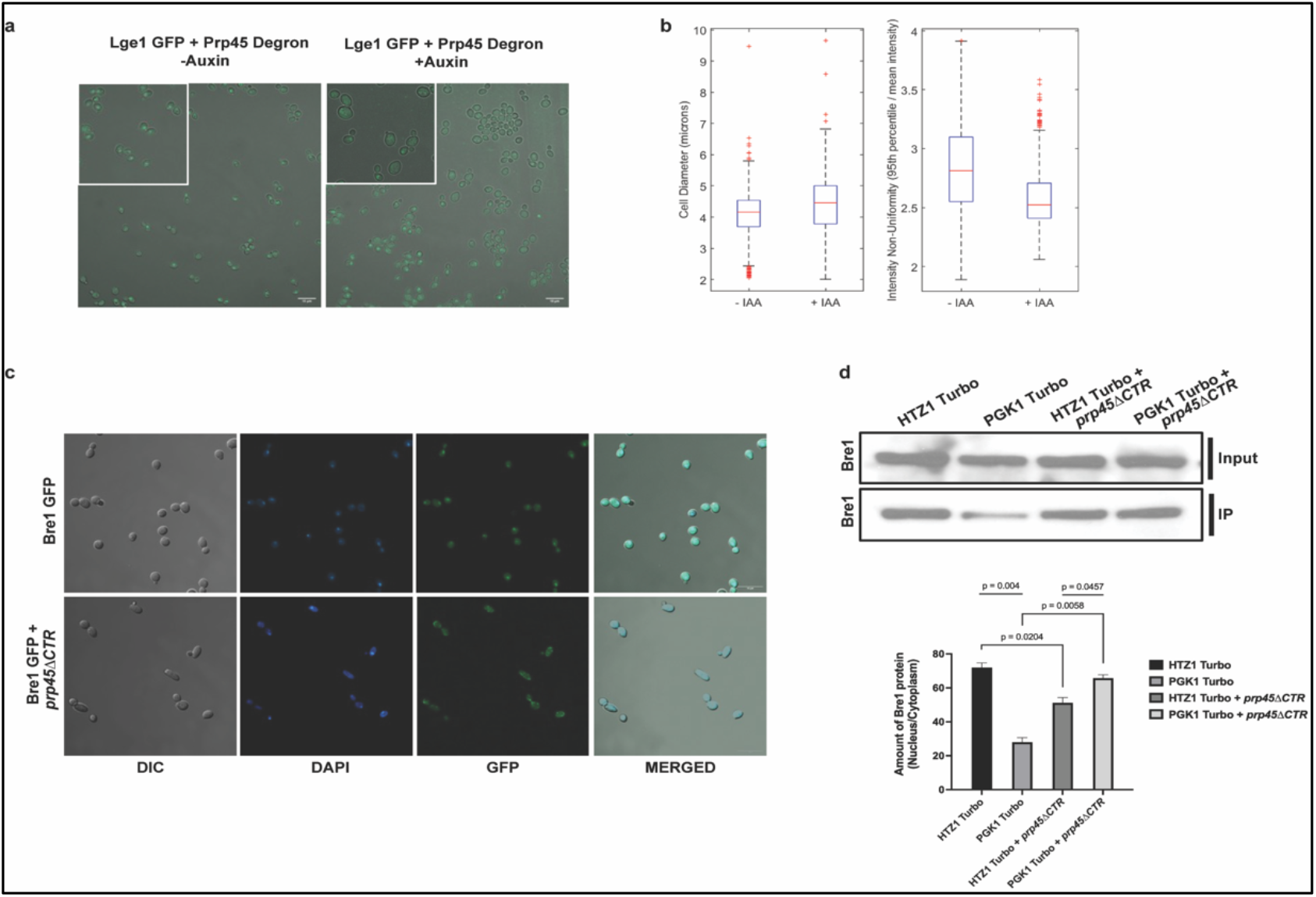
Auxin-mediated Prp45 degradation causes Lge1 degradation, and altered distribution of Bre1 between the nucleus and cytoplasm. (a) Reduction in Lge1 intensity and puncta formation in the nucleus. The intensity of Lge1-GFP was measured using confocal microscopy at 63X in a strain in which Prp45 was degraded with auxin. (b) Cell segmentation and intensity quantification. Confocal images of cells treated with or without auxin were segmented to measure cell size and GFP intensity using CellPose. Cell size and GFP intensity were quantified and plotted in the graph. (c) Distribution of Bre1 is altered between the nucleus and cytoplasm in the *prp45ΔCTR* strain. Bre1 was GFP tagged in WT and *prp45ΔCTR*. Confocal images show the altered distribution of GFP-tagged Bre1 in the *prp45ΔCTR* strain. (d) Protein transit assay. Immunoblot showing a change in biotinylated Bre1 protein, either by HTZ1-Turbo in the nucleus or PGK1-Turbo in the cytoplasm. The data were quantified and plotted using GraphPad PRISM.

To determine how this decrease in Lge1 and Lge1-containing puncta affects Bre1 activity, Bre1 was GFP-tagged in wild-type and *prp45ΔCTR* cells. While Bre1 is highly concentrated in the nucleus in WT cells, it is evenly distributed between the nucleus and cytoplasm in *prp45 ΔCTR* cells (Fig. 5c). We further confirmed this observation using a protein transit assay in which the nuclear histone protein H2A.Z or the cytoplasmic protein Pgk1 was turboID (Biotin ligase or Bir A) tagged in cells with either WT Prp45 or Prp45 with a deleted C-terminal end. The cells were grown in the presence of biotin, and the biotinylated proteins were pulled down with streptavidin beads. Bre1 was detected using an anti-GFP antibody. If Bre1 protein is largely chromatin-bound, as we suspect is the case in wild-type cells, then more Bre1 will be biotinylated in the Htz1-turboID cells and less will be biotinylated in the cytoplasmic Pgk1 TurboID cells (Fig. 5d & graph). If the distribution is altered as in *prp45ΔCTR* cells, we expect to see more Bre1 biotinylated in the cytoplasm by PGK1 TurboID relative to the levels observed in WT cells, which is precisely what is observed (Fig. 5d & graph).

As shown in figure 1, the highly disordered domain of Prp45 extends to the conserved SH2 domain of the protein, and the region of the protein up to the SH2 domain that includes the disordered region is sufficient to confer near WT growth (Fig. 1). To clarify that, indeed, the entire disordered portion of the C-terminus is necessary and sufficient to confer the H2B ubiquitination functions, a Prp45 truncation containing the disordered region of the C-terminus, *prp45 (1-260),* was analyzed for each of the phenotypes associated with *prp45ΔCTR*. Lge1 stability (Fig. S5b), H2B ubiquitination (Fig. S5c), and cell size homeostasis (Fig. S5d). As with growth, *prp45 (1-260)* is sufficient to restore near-WT activity. However, not only does 1-184 show poor growth, but Lge1 is unstable in this mutant, consistent with a requirement for the entire IDR for Lge1 stability (Fig. S5b). Finally, the protein level of Prp45 (1-260) was also checked (Fig. 5a) suggesting that the protein is expressed normally.

### The protective C-terminal IDR added in trans is sufficient for the chromatin function and is functionally conserved from yeast to humans

The C-terminal portion of Prp45, containing the disordered domain, is critical for Prp45 function in chromatin modification. This region of Prp45 also contributes to its stability and to its ability to interact with and modulate the activity of the H2B ubiquitination machinery. Given these findings, we considered the possibility that the C-terminus could act in trans with the N-terminal part of the protein to affect Prp45 function. To test this, the C-terminal domain of Prp45 (170-379) was cloned into a yeast 2µ plasmid under the native Prp45 promoter and expressed in the *prp45ΔCTR* strain. Intriguingly, the C-terminus expressed in trans rescues the growth defect (Fig. 6a and Fig. S6a). To determine if the C-terminus expressed in trans can rescue the chromatin-related phenotypes, Lge1 stability was assessed when the C-terminal domain (170-379) was expressed in the *prp45ΔCTR* strain. Trans expression of the C-terminal domain stabilizes the Lge1 protein in the *prp45ΔCTR* strain (Fig. S6b upper image). Moreover, the trans-expressed C-terminal domain in *Kan^r^ Lge1 CDS degron prp45ΔCTR* strain partially restores growth on IAA + Kan plate, indicating that Lge1 becomes more stable and less susceptible to proteasome-mediated degradation in the presence of the C-terminus (Fig. S6b growth assay, lower image). Since trans expression of the C-terminal domain stabilizes the Lge1 protein, we predicted that it would also rescue H2B ubiquitination. Indeed, H2B ubiquitination and related phenotypes are restored (Fig. S6c).

**Figure 6.**
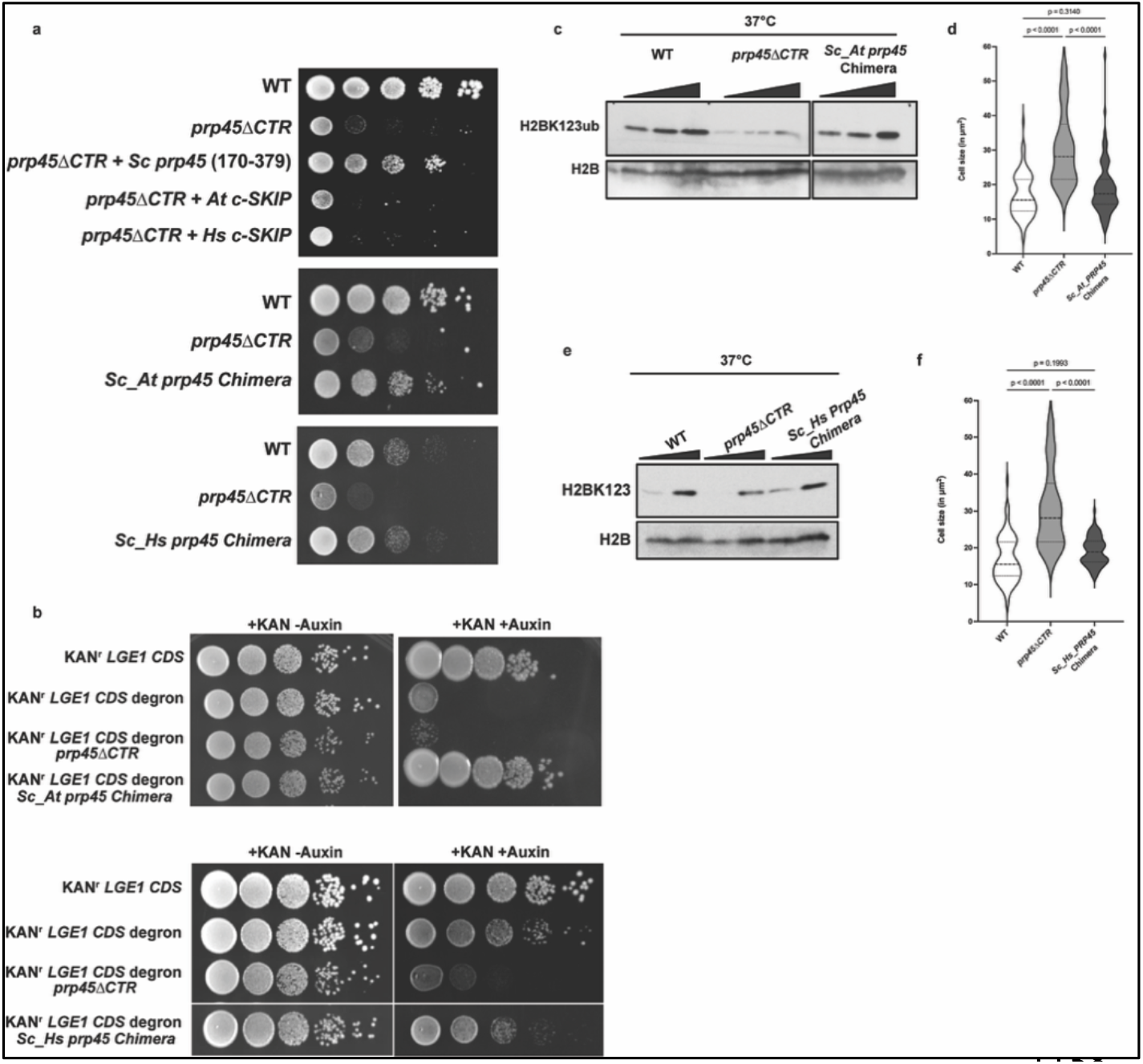
The function of Prp45’s C-terminal domain is evolutionary conserved. (a) (Top panel) Trans expression of the C-terminal domain rescues the growth defect at high temperature. WT, *prp45ΔCTR*, *prp45ΔCTR* + (170-379) of Prp45, expressed from a plasmid (pRS315) were grown on plates containing Sc-Leu media. The C-terminal region of the *Arabidopsis thaliana* SKIP protein and the C-terminal region of SKIP from *Homo sapiens* (Fibroblast cells) was expressed from pRS315 plasmid in the *prp45ΔCTR* strain and compared to the *prp45ΔCTR* strain. Cells were serially diluted and incubated at 37°C for six days. The fusion of *At-*SKIP and the *Sc* Prp45 N-terminus (1-169) (middle panel) or *Hs*-SKIP and the *Sc* Prp45 N-terminus (1-169) (lower panel), were grown on plates containing WT cells and prp45ΔCTR. (b) Growth assay showing Lge1 stability in the *Arabidopsis* and Human Prp45 chimera strains. Orthogonal degron assay described in Fig. 3d was done. (c) The Sc_At chimera can restore H2B monoubiquitination. An Immunoblot showing H2BK123 ubiquitination and H2B from whole cell extracts from WT, *prp45ΔCTR,* and *Sc_At PRP45* chimera strains. (d) The chimera restores yeast cell size. WT, *prp45ΔCTR,* and *Sc_At PRP45* chimera strains grown to OD_600_ 0.6 at 37°C. Confocal imaging (DIC) was performed, and cell size was measured using ImageJ. Cell size was plotted in the graph using GraphPad PRISM, and an unpaired one-way ANOVA with Tukey’s post hoc analysis (n = 80) was used to calculate the *p*-values. (e) Growth assay showing Lge1 stability in the human chimera strain. Lge1 was N and C-terminally tagged with Kan^r^ and degron in wild type, *prp45ΔCTR* and the *Sc_Hs PRP45* chimera strains. Cells were tenfold serially diluted on a YPD + kanamycin + IAA plate and incubated at 30°C. (f) The *Sc_Hs* chimera can also restore H2B monoubiquitination. An Immunoblot showing H2BK123 ubiquitination and H2B from whole cell extracts isolated from WT, *prp45ΔCTR,* and *Sc_Hs PRP45* chimera strains (g) The Human chimera restores yeast cell size. WT, *prp45ΔCTR,* and *Sc_Hs PRP45* chimera strains grown to OD_600_ 0.6 at 37°C. Confocal imaging (DIC) and cell size was measured as mentioned before followed by graph plotting and statistical analysis.

The observation that the two halves of Prp45 can function on separate molecules suggests that the N- and C-termini of the protein may make functional interactions. Data from both Dictyostelium and *S. Pombe* Prp45 show strong evidence of Prp45-Prp45 interaction (dimerization).^46^ Furthermore, when Prp45 is analyzed using AlphaFold, not only is dimerization predicted, but the modeling also predicts interactions between the N- and C-termini of the proteins.

To determine if the C-terminus of Prp45 can function independently of the N-terminus, we expressed just the C-terminal portion of the protein alone to determine if it could suppress the lethal phenotypes associated with the Prp45 deletion. *S*trains containing only the C-terminal domains (170-379, 175-379, 184-296) after the WT plasmid was “shuffled out” were analyzed for growth on 5-FOA. The C-terminus alone does not restore *prp45Δ* growth on SC-FOA plates (Fig. S6d).

The fact that the C-terminal domain, expressed in trans, can partially restore growth and H2B ubiquitination highlights the importance of this disordered region. As shown in figure 1B, this feature of the protein is conserved, down to the size of the predicted region of helical propensity, suggesting that homologs of Prp45 may also stabilize interactions that drive H2B ubiquitination. To test this, the C-terminal domains from *Arabidopsis* (*At* C-SKIP) and Human fibroblast (*Hs* C-SKIP) were cloned into a yeast 2µ plasmid to determine if they could restore Prp45 functions in trans. The C-terminal regions from *At* and *Hs* alone do not rescue the growth defect (Fig. 6a upper image). However, when At SKIP is fused to the N-terminus of the yeast protein, *Sc_At prp45 Chimera* [*Sc* Prp45 (1-169) + *At* (C-terminal region], strong rescue of the growth defect is observed (Fig. 6a middle image), suggesting that constraining this domain to be within proximity of the N-terminus is critical for growth rescue. Similarly, an *Sc_Hs* prp45 chimera also restores growth to the *prp45 ΔCTR* cells (Fig. 6a, lower image).

Previous studies of Prp45 report that overexpression of either the *Arabidopsis* or human SKIP protein can replace the yeast protein.^19^ However, when either of these proteins is expressed from the yeast 2µ plasmid in the absence of Prp45 (*prp45Δ* strain), little 37°C growth rescue is observed (Fig. S6d). On the other hand, when either *At* or *Hs* SKIP in the yeast 2µ plasmid is expressed in *the prp45ΔCTR* strain, where the N-terminal portion of the protein is present, the full-length SKIP proteins can rescue the growth defect (Fig. S6a), with the A*t* SKIP conferring better rescue. Together, these data suggest an evolutionarily conserved function for the C-terminus of Prp45/SKIP and support a model whereby protein interaction between the C- and N-termini (such as through dimerization of the proteins) is important for their functions.

The observation that the chimera protein can support growth suggests that these protein interactions also form a surface that supports Prp45 interaction with and stabilization of Lge1, as well as proper H2B ubiquitination. Lge1 stability was evaluated as described in Figure 3d, in which the stability of the protein was assessed by growth on the IAA + Kanamycin plate. The cells grow well, indicating that the chimeric protein protects Lge1 from degradation (Fig. 6b). Moreover, H2B ubiquitination is also restored when the IDR of Prp45 from *Arabidopsis* is fused to Prp45ΔCTR (Fig. 6c). With the restoration of H2B ubiquitination, we predicted that aberrant cell morphology and size would also be restored by the chimera. Indeed, this is what we observe (Fig. 6d). The C-terminal portion of the Human SKIP protein behaves similarly when fused to Prp45ΔCTR. A chimeric protein comprised of the C-terminal half of Human SKIP and the N-terminus of *Sc* prp45 (*Sc_Hs* prp45 chimera) not only restores growth (Fig. 6a lower growth assay), but also Lge1 stability (Fig. 6b lower growth assay), H2B monoubiquitination (Fig. 6e), and cell size (Fig. 6f). The results show that the mechanism whereby Prp45 affects chromatin modification is encoded in the conserved structural feature of the protein rather than its sequence, a feature that has persisted across millions of years of evolution.

The observation that the N-terminus of the yeast protein interacting with either the *Arabidopsis* or human SKIP protein supports growth (Fig. S6a) suggests that these interactions form a surface critical for SKIP activity in H2B ubiquitination. Intriguingly, SKIP from *Arabidopsis* is critical for flowering and Histone H2BK120 monoubiquitination,^47^ consistent with an evolutionarily conserved role for Prp45/SKIP in H2B ubiquitination. To test whether the full-length SKIP proteins support conditions necessary for proper H2B ubiquitination, we analyzed Lge1 stability, H2B ubiquitination, and H2B-dependent cell morphology when either *At* SKIP or *Hs* SKIP is expressed in *prp45ΔCTR* cells. Indeed, Lge1 protein becomes stable in *prp45ΔCTR* cells expressing A*t* SKIP or *Hs* SKIP (Fig. S6e). H2B ubiquitination (Fig. S6f and S6g) and cell size are also restored (Fig. S6h). Although SKIP proteins from metazoans long diverged from *S. cerevisiae*, it appears that the role of the disordered region of the protein, the C-terminus, in protein interactions that are critical for Prp45/SKIP activity has been conserved.

## Discussion

Prp45 is structurally conserved from yeast to humans, and while the *S. cerevisiae* protein has been characterized for its role in splicing, its homologs in *Arabidopsis thaliana* and mammals play roles in transcriptional events involved in flowering time initiation and transcriptional activation, respectively.^18,19^ Here we show that Prp45 helps to direct H2B monoubiquitination through its C-terminal, intrinsically disordered region, by a mechanism distinct from its established role in splicing fidelity. This work therefore provides a mechanism for a direct role for a splicing factor in chromatin modification (Fig. 7). Rather than simply providing passive structural support, the Prp45 C-terminus protects Lge1 from active turnover. In its absence, Lge1 is degraded through the SUMO-targeted ubiquitin ligase Slx5, and *SLX5* deletion restores Lge1, H2B ubiquitination, cell morphology, and growth.

**Figure 7.**
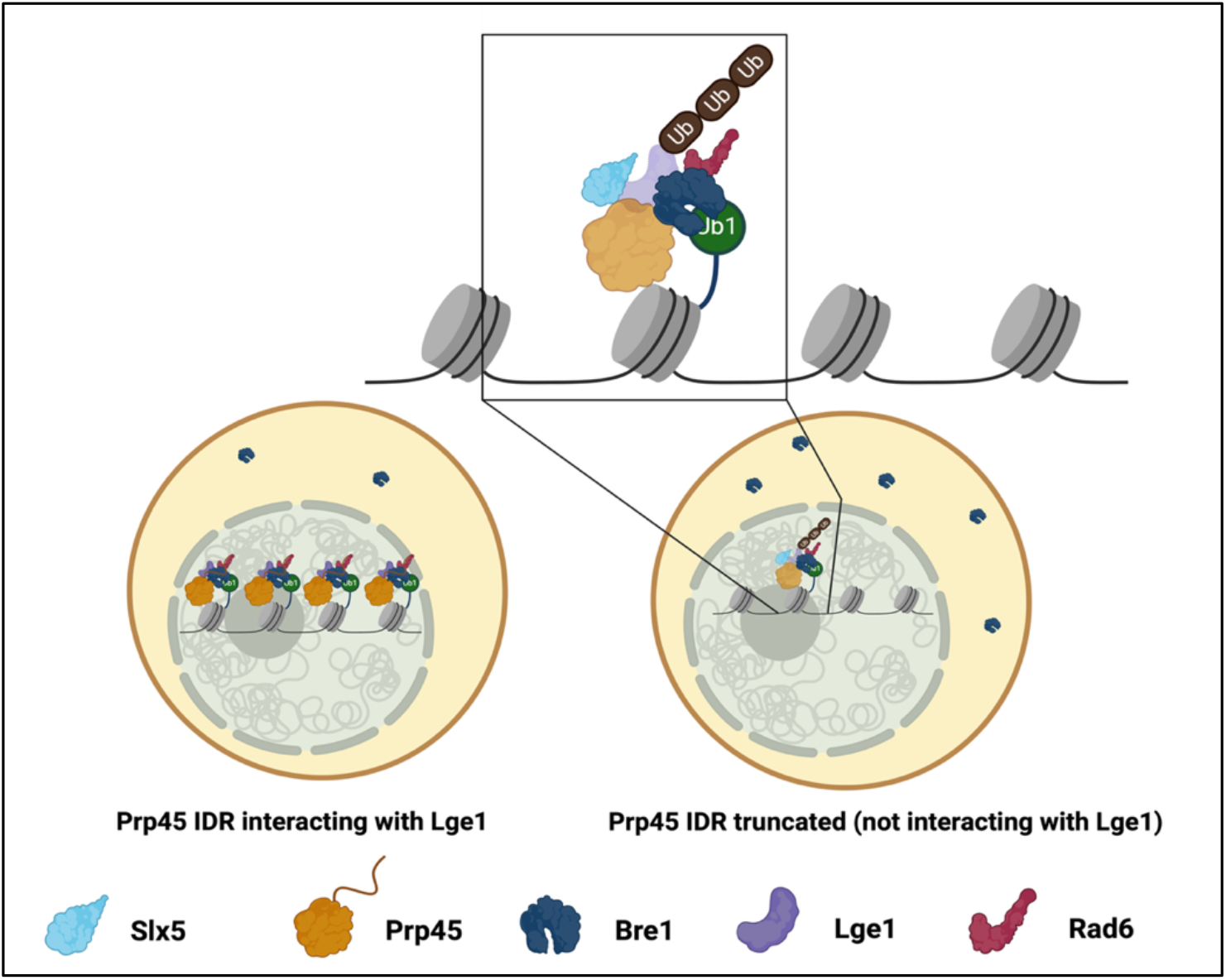
Model of Prp45 effects on histone H2B ubiquitination. The model shows that truncation of the C-terminal domain (orange) renders the Lge1 protein (light blue) unstable. In wild-type cells (Left), the Prp45 C-terminus protects Lge1 from the SUMO-targeted ubiquitin ligase Slx5. Disruption of the Prp45 CTR-Lge1 interaction, such as by CTR truncation (Right), exposes Lge1 to Slx5-dependent turnover and decreased Lge1 stability. The biomolecular condensate formed by the Lge1 protein is also significantly reduced, and Lge1 can no longer concentrate Bre1 (dark blue) in the nucleus. Therefore, H2B ubiquitination is abrogated. Model was generated using BioRender.

These findings place the Prp45–Lge1 interaction at a node of crosstalk between the ubiquitin and SUMO systems. Slx5 couples SUMO modification to ubiquitin-dependent degradation; our data indicate that the abundance of Lge1, and therefore the output of the Bre1 H2B-ubiquitination machinery, is determined by the balance between Prp45-C-terminus-mediated protection and Slx5-directed turnover. In this view, the intrinsically disordered region does not merely tether Prp45 to Lge1; it modulates Lge1’s exposure to protein quality control. Because Lge1 forms puncta that locally concentrate Bre1, and because STUbL pathways are increasingly recognized as regulators of condensate composition and turnover,^48^ Slx5 may govern H2B ubiquitination by controlling the stability of a phase-separating scaffold. An important open question is whether Lge1 is itself SUMO-modified and whether such modification is antagonized by the Prp45 C-terminus.

These studies highlight the importance of low-complexity, disordered regions more generally. Although about 50 aa of the Prp45 C-terminus have the propensity to form an alpha-helical structure, which is likely affected by protein interactions, much of the protein is highly disordered, with an extensive low-complexity domain (LCD). LCDs facilitate the formation of biomolecular condensates,^49,50^ thus separating specific biochemical functions and coordinating biochemical processes.^51,52,53^ Lge1 is another striking example: approximately 85% of its residues lie in a disordered region (Fig. S4a upper image), and this disorder is critical for Bre1’s recruitment to nucleosomes and H2B monoubiquitination. Consistent with a role for the Prp45 IDR in sustaining this assembly, loss of the C-terminus reduces Lge1 puncta and disperses Bre1 from the nucleus.

Trans expression of the C-terminal domain restores growth, cell size, and H2B monoubiquitination, further supporting a critical role for this region. Remarkably, yeast and metazoan Prp45 proteins share no sequence similarity beyond the evolutionarily conserved SNWKN motif (Fig. S1), which is essential for Prp45 interaction with the spliceosome. Yet, while the C-terminal domains from *Arabidopsis* and human cells alone cannot rescue the growth defect, fusing them to the *S. cerevisiae* N-terminal domain rescues growth, H2B monoubiquitination, and cell size, indicating evolutionary conservation of a disordered, sequence-independent regulatory function. WAC (WW domain-containing adaptor with coiled-coil), the Lge1 homolog in mammals, likewise contains an extensive N-terminal disordered region and a conserved coiled-coil domain;^38^ it interacts through that coiled-coil with RNF20/40, the mammalian homolog of Bre1, while its N-terminal WW-containing domain binds RNA polymerase II and recruits RNF20/40 for H2B monoubiquitination and transcriptional regulation. H2B monoubiquitination is completely lost upon WAC deletion.^54^ In plants, the Bre1 homologs HUB1 and HUB2 are responsible for H2B monoubiquitination, but no Lge1 homolog has been found.^55^

The Prp45 C-terminus–Lge1 interaction has clear physiological consequences. *prp45ΔCTR* cells show a significant reduction in H2B monoubiquitination and become large and elongated, a morphology that is reversed by deleting *UBP8*. The same enlarged-cell phenotype follows loss of H2B monoubiquitination in *lge1Δ* or *bre1Δ* cells,^36^ and in mammalian cells H2B monoubiquitination is required for effective activation of cell cycle checkpoints during genotoxic stress.^54^ Intriguingly, Abramova et al. report inefficient splicing of some cell cycle regulators, such as *ACT1*, when Prp45 is truncated. ^56^ Together these observations suggest that the combined effect of splicing and H2B monoubiquitination underlies the morphological defect.

Our genetic data as well as the proximity labeling experiments indicate that the C-terminus does more than this one job. While C-terminal truncation of Prp45 destabilizes Lge1, combining *prp45ΔCTR* with *lge1Δ* causes a substantially more severe defect, as does combining it with *bre1Δ*. Deletion of *BRE1* or *LGE1* results in a complete loss of H2B monoubiquitination, and when these deletions are combined with *prp45ΔCTR* the outcome is synthetic lethality (Fig. 2a). This epistasis is consistent with the C-terminal domain having functions in addition to its now-established role in H2B monoubiquitination, such as RNA splicing.

An outstanding question is how the interactions described here affect intron-containing genes. H2B monoubiquitination is enriched at highly expressed genes, particularly the intron-containing ribosomal protein genes,^57^ and it marks exon–intron boundaries in both yeast and human genomes,^22,58,59^ where the splicing of a subset of transcripts is particularly sensitive to this modification. Chromatin has long been understood to shape spliceosome assembly and splicing outcomes. Here, we show that this relationship also occurs in reverse, with a spliceosomal protein determining, through a disordered region, the fate of the machinery that marks a key chromatin modification. The two processes are therefore not merely coexisting in time and space, but appear to be reciprocally coupled, and how a cell keeps this loop in balance is the question these findings open. The results with the chimeric proteins demonstrate that this mechanism is conserved not a sequence but a capacity. This capacity lies in the ability of a disordered region to engage in protein-protein interactions such that a partner protein is shielded from the activity of the degradation machinery. It is important to test hypothesis that the disordered region in metazoan SKIP affects the stability or activity of the H2B ubiquitination machinery—a prediction that is now directly testable.

## Supporting information

Supplemental figures, tables, strains and primer list

## Acknowledgements

AP, TLJ, and the work in the Johnson lab are supported by the NIH (R35GM149290).

We would like to express our gratitude to Dr. Hillary Coller for providing Human Fibroblast cells, and Dr. Jeffrey A. Long for providing *Arabidopsis* leaves for RNA isolation. Thank you also to Drs. Long and Pavak Shah for access to confocal microscopy facilities and for assistance with confocal image analysis. Thank you to Dr. Azad Hossain and members of the Johnson Lab for careful reading of the manuscript.

The authors would like to acknowledge the UCLA Core Mass Spectrometry Facility for their technical support and access to instrumentation.

## Materials and Methods

### Strains and growth conditions

All strains of *S. cerevisiae* used in this study are listed in Table S1. All strains are derived from BY4741. The Prp45 degron strain was generated by integrating Adh1/AtTIR1 into the HIS3 locus. AID-HA-Hyg was added to the C-terminal of endogenous Prp45 by standard replacement with a marker cassette.^60^ Endogenous C-terminal tagged strains were generated by homologous recombination followed by PCR amplification from the pFA6a-3HA-kanMX6 and pFA6a-5FLAG-NatMX6 plasmids as described previously.^60^ *Lge1*, *UBP8*, *UBP10*, and *Bre1* were knocked out by replacing the KanMX6 cassette. The C-terminal domain (amino acids 170-379, 175-379, and 184-296) of Prp45, along with the promoter sequence, was PCR-amplified with CloneAmp (Takara) using primers listed in the supplementary material. PCR was performed for 35 cycles, with denaturation at 98°C for 15 seconds, extension at 55°C for 15 seconds, and a final extension at 72°C for 20 seconds. The C-terminal domain or the full length of *At* SKIP and *Hs* SKIP were cloned under *Sc* Prp45’s promoter sequence. Amplicon was run on 1.8% agarose gel and gel-purified using the Qiagen gel extraction kit. The purified amplicon was subjected to restriction digestion with *BamHI* and *SalI* and cloned in *BamHI* and *SalI* restriction sites in pRS315 vector (Addgene). The chimeric Prp45 was generated by using primers used to amplify the C-terminal region of *At* SKIP, and the homologous recombination technique was used to integrate it into the yeast genome. Cells were grown on YPD (1% yeast extract, 2% peptone, and 2% glucose) or synthetic dropout media at 30°C and 37°C.

### Growth assay

5mL overnight cultures were transferred to a fresh 50 mL YPD or selective dropout media (SC-URA or SC-Leu) and grown at 30°C until the OD reached 0.5. Cells were tenfold serially diluted and spotted on YPD or YPD + Kan + IAA or SC-Leu or SC FOA-URA plates. Plates were kept at 30°C and 37°C to observe the effect of elevated temperature.

### Proximity labeling, biotinylation, and Mass-spectrometry

5 mL TurboID-tagged (Biotin ligase) overnight cultures were transferred to fresh 50 mL YPD supplemented with 50 µM biotin. Cells were grown at 30°C until the cultures reached an OD of 0.8. Cell pellets were resuspended in 250 μL of RIPA buffer (50 mM Tris-HCl, pH 7.5, 150 mM NaCl, 1.5 mM MgCl_2_, 1 mM EGTA, 0.1% SDS, 1% NP-40, supplemented with 0.4% sodium deoxycholate, 1 mM DTT, 1 mM PMSF, and 1x complete). Acid-washed beads were added to the resuspension, and samples were lysed in a cell disruptor for 10 minutes at 4°C. The lysates were centrifuged at 13,000 rpm for 15 minutes at 4 °C to remove insoluble material. Lysates were briefly sonicated and treated with benzonase (Medchem exp) for 3 hours at 4°C to digest DNA and RNA. 10 mg of total proteins were subjected to affinity purification using 50 μL streptavidin (GE) at 4°C for 3 hours. Beads were then washed once with wash buffer (50 mM Tris-HCl pH 7.5, 2% SDS) followed by three times wash with RIPA buffer containing DTT (50 mM Tris-HCl pH 7.5, 150 mM NaCl, 1.5 mM MgCl_2_, 1 mM EGTA, 0.1% SDS, 1% NP-40, 1 mM DTT). Finally, the beads were washed five times with 20 mM ammonium bicarbonate. All the washes were done for 5 minutes at room temperature in a nutator. For mass spectrometric analysis, streptavidin-bound proteins were reduced with 10 mM DTT, followed by alkylation with 15 mM iodoacetamide.

Protein-bound beads were subjected to trypsin digestion at 37°C overnight, then stopped by the addition of formic acid (final concentration 1%). Peptides were then extracted with acetonitrile, lyophilized, and resuspended in 0.1% trifluoroacetic acid (TFA) for desalting on ZipTips. (EMD Millipore) The cleanup of peptide samples was performed as described previously.^61^ Trypsin-digested samples were analyzed by LC/MS. The MaxQuant software package version 1.5.1.2 was used to analyze the data against the *S. cerevisiae* UniProt proteome. The peak intensity of peptides was used for relative quantification of proteins.

### *In vivo* ubiquitination assay

25 mL of overnight saturated cultures (Wild type + YEp181 CUP1 His Ub and *prp45ΔCTR* + YEp181 CUP1 His Ub) were transferred to 200 mL Sc-Leu media and allowed to grow at 30°C until they reached an OD of 0.6. For the induction of histidine tagged ubiquitin 500 μM of CuSO_4_ was added in each culture and then the cultures were transferred to 37°C for 3 hours. Cells pelleted and washed with ice cold water and then resuspended in 3mL of Denaturing lysis buffer (6M Guanidine-HCl, 100 mM Sodium phosphate buffer, 10 mM Tris-Cl pH-8.0, 300 mM NaCl, 1% Triton X-100, 20 mM N-Ethyl maleimide, 10 mM imidazole and protease inhibitor). An equal volume of 0.5 mm acid-washed glass beads (BioSpec) was added to the cell resuspension, and the samples were lysed in a cell disruptor for 10 minutes at room temperature. Samples were spun at 3000g for 5 minutes and supernatants were collected and transferred to 1.5 mL microcentrifuge tube. Lysates were briefly sonicated (10 sec on 20 sec OFF for 3 cycles at 30% amplitude) and spun at 16000g for 15 minutes. Supernatants were collected and transferred to a 15 mL conical tube, 100 uL of pre-calibrated Ni-NTA agarose beads were added and the conical tubes were placed on a nutator for 2 hours at room temperature.

Beads were washed 3 times with the denaturing lysis buffer followed by 2 times wash with lysis buffer containing 20 mM imidazole. Beads were washed again with wash buffer containing 8M urea, 100 mM Sodium phosphate buffer, 100 mM tris-cl pH-6.3 and 20 mM imidazole. After the final wash his tagged ubiquitinated proteins were eluted using elution buffer (8M urea, 250 mM imidazole). Proteins were precipitated with 100% ice cold acetone for 30 minutes at −20°C followed by spin at 16000g for 15 minutes at 4°C. Pellets were washed 3 times with 80% ice cold acetone followed by a brief drying in a desiccators. Finally, the pellets were dissolved in 50 μL of 1X SDS PAGE buffer.

### Microscopy and image quantification

Wild type and mutant strains were stained with 5 μM DAPI and observed under the 63X oil immersion lens. The images were quantified using ImageJ. Cell counting and segmentation was done using cell pose. In house MATLAB code was used to analyze and quantify the images.

### Statistical analysis

All experiments were performed a minimum of three times. Statistics in this study were presented as mean ± SD. Error bars represented SD in triplicate experiments. Statistical significance (*p*-values) for comparison was assessed by One-way ANOVA with Tukey post-hoc analysis.

### In vivo depletion of Prp45 and Lge1

Prp45-AID-HA and Lge1 AID-HA strains were grown on YPD at 30°C for 1.5 hours, followed by 1 mM IAA (I5148, sigma) treatment for different time points. Cells were spun, and the pellets were kept at −80 °C.

### Yeast nuclear extract preparation and Immunoblotting

Saturated overnight 5 mL cultures were transferred to 50 mL and allowed to grow at 30°C until they reached an OD of 0.8-1. For elevated-temperature stress, cells were first grown at 30°C for 1 hour, then transferred to 37°C and grown for 2 hours. Cells were pelleted and resuspended in 500μL of lysis buffer (50 mM HEPES, pH 7.5, 150 mM NaCl, 1 mM EDTA, 1% Triton X-100, 0.1% sodium deoxycholate) with protease inhibitor. An equal volume of 0.5 mm acid-washed glass beads (BioSpec) was added to the cell resuspension, and the samples were lysed in a cell disruptor for 10 minutes at 4°C followed by ultracentrifugation at 15,000g for 15 minutes. Lysates were clarified and aliquoted and frozen liquid nitrogen. Proteins were separated in an 8% SDS-PAGE for Lge1-FLAG, Prp45AID-HA, and a 15% SDS-PAGE for histone H2BK123 monoubiquitination and histone H2B. HA blots were probed with anti-HA antibody (clone 12CA5, Roche), FLAG blots were probed with anti-FLAG antibody (Sigma, catalog no F1804), histone H2BK123Ub1 blots were probed with anti-H2BK123Ub1 (anti-ubiquityl H2B,5546S; CST, Cell Signaling Technology, Danvers, MA, USA), and histone H2B blots were probed with anti-H2B (Active motif, catalog no 39210). Immunoblots were quantified using ImageJ software.

### Co-immunoprecipitation

25 mL overnight saturated culture was transferred to 125 mL of fresh YPD, and cells were grown until A_600_=0.8-1. Cells were then pelleted and resuspended in 1 mL of lysis buffer with protease inhibitors. Acid-washed beads were added to the resuspension, and samples were lysed in a cell disruptor for 10 minutes (one minute ON, one minute OFF) at 4°C. The lysates were centrifuged at 13,000 rpm for 15 minutes at 4 °C to remove insoluble material.

1 mL of lysate was precleared with 50 μL of Protein A/G agarose beads (sc-2003, Santa Cruz Biotechnology). 25 μL of precleared lysate was removed as input. 4 μg of anti-HA antibody (clone: 12CA5, Roche) was added to the 700 μL of clarified lysate and rotated at 4°C overnight. Samples were incubated with 50 µL of Protein A/G agarose beads for 2 hours at 4°C. Beads were then washed five times with lysis buffer and eluted by boiling in 2X SDS-PAGE loading buffer.

### RNA isolation

Yeast cells were grown until the *A*_600_ = 1.0. Total RNA was isolated using the hot acid phenol method. Yeast cells were treated with acidic phenol (P4682; Sigma) at 65 °C, and the RNA was extracted with phenol/chloroform/isoamyl alcohol with SDS, followed by ethanol precipitation. Following total RNA isolation, 25 μg of RNA was treated with DNase I (Roche). 3 μg of DNase I-treated RNA was used for cDNA synthesis. The first strand of cDNA was generated using Maxima Reverse Transcriptase (ThermoFisher Scientific). Gene-specific primers are listed in the supplementary material table S1. 1–2 µL of cDNA was used as the template in a 25 µL PCR reaction.

### *In silico* protein interaction analysis

Alphafold2 was used to generate three-dimensional protein structures, and Google Colabfold was used to analyze the protein-protein interaction. The best fit model was fetched to ChimeraX 1.9, and subsequent analysis was done in ChimeraX 1.9, including probable contact sites between the interacting proteins.

### Cycloheximide chase experiment

5 mL overnight saturated cultures were transferred to 50 mL and allowed to grow at 30°C until the OD of 0.6 is reached. Cells were then transferred to 37°C for 1 hour and then cycloheximide was added at a concentration of 100 μg/mL to the cultures. Cells were collected and pelleted at different time points up to 2 hours, washed with ice cold water and flash frozen in liquid nitrogen followed by protein isolation that has been mentioned before.

### Modified Protein transit assay

Nuclear protein H2A.Z (encoded by *HTZ1*) and cytoplasmic protein (PGK1) were C-terminally tagged with Turbo ID (biotin ligase) by homologous recombination, followed by PCR amplification from the pFA6a-Turbo-ID-3 Myc-kanMX6. 5 mL TurboID-tagged overnight cultures were transferred to fresh 50mL YPD supplemented with 50 µM biotin. Cells were grown at 30°C until the cultures reached an OD of 0.8. Cell pellets were resuspended in 250μL of RIPA buffer (50 mM Tris-HCl, pH 7.5, 150 mM NaCl, 1.5 mM MgCl_2_, 1 mM EGTA, 0.1% SDS, 1% NP-40, supplemented with 0.4% sodium deoxycholate, 1 mM DTT, 1 mM PMSF, and 1x complete). Acid-washed beads were added to the resuspension, and samples were lysed in a cell disruptor for 10 minutes at 4°C. The lysates were centrifuged at 13,000 rpm for 15 minutes at 4 °C to remove insoluble material. Lysates were briefly sonicated and treated with benzonase (Medchem exp) for 3 hours at 4°C to digest DNA and RNA. 30mg of total proteins were subjected to affinity purification using 50 μL streptavidin (GE) at 4°C for 3 hours. Beads were then washed once with wash buffer (50 mM Tris-HCl pH 7.5, 2% SDS) followed by three times wash with RIPA buffer containing DTT (50 mM Tris-HCl pH 7.5, 150 mM NaCl, 1.5 mM MgCl_2_, 1 mM EGTA, 0.1% SDS, 1% NP-40, 1 mM DTT). Finally, the beads were washed five times with 20 mM ammonium bicarbonate. All the washes were done for 5 minutes at room temperature in a nutator. Beads were then washed twice with lysis buffer and half of the beads were sent to mass spectrometric analysis and other half were eluted by boiling in 2X SDS-PAGE loading buffer.

