## Supplemental figures, tables, strains and primer list for "A conserved disordered region of a splicing factor promotes histone H2B ubiquitination by opposing SUMO-targeted degradation"

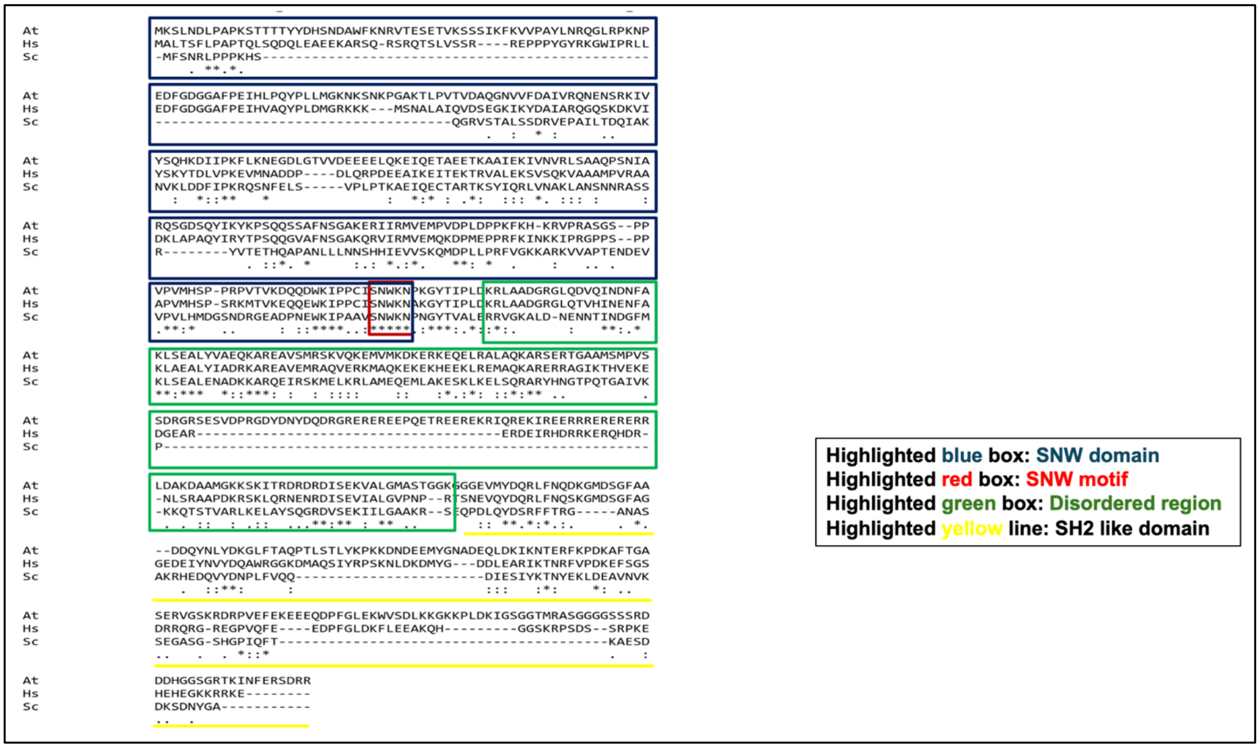


**Supplementary Figure 1**. **Prp45 and its homolog's Sequence alignment.**

(a) Sequence alignment. UniProt IDs were used to retrieve the sequences of Prp45 from *S. cerevisiae*, *Arabidopsis thaliana*, and *Homo sapiens*; sequence alignment was performed using ClustalW.

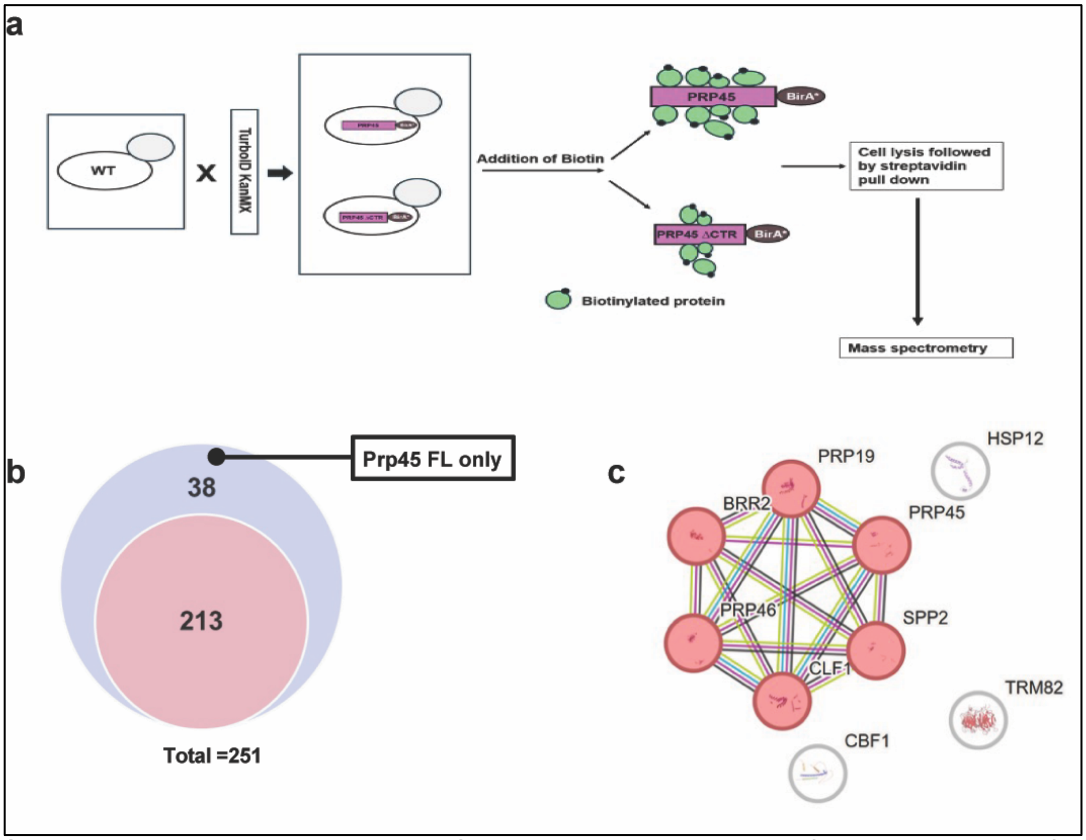


**Supplementary Figure 2. The C-terminal region-specific interactors of Prp45 identified by in vivo biotinylation and loss of the C-terminal domain also affect its stability.**

(a) Schematic of the *in vivo* biotinylation assay, where full-length Prp45 and Prp45ΔCTR are C-terminally Biotin ligase tagged. Model was generated using BioRender. (b) Interactors identified from mass spectrometry data. The Venn diagram indicates 38 unique interactors specific to the full-length Prp45 only and 213 interactions that occur with both. (c) Hierarchical clustering of the interactors that show a significant change in interaction with Prp45 FL and Prp45ΔCTR. BRR2, SPP2, PRP46, CLF1, PRP45, CBF1, TRM82, and HSP12 show a significant reduction in interaction when the C-terminal end of Prp45 is deleted.

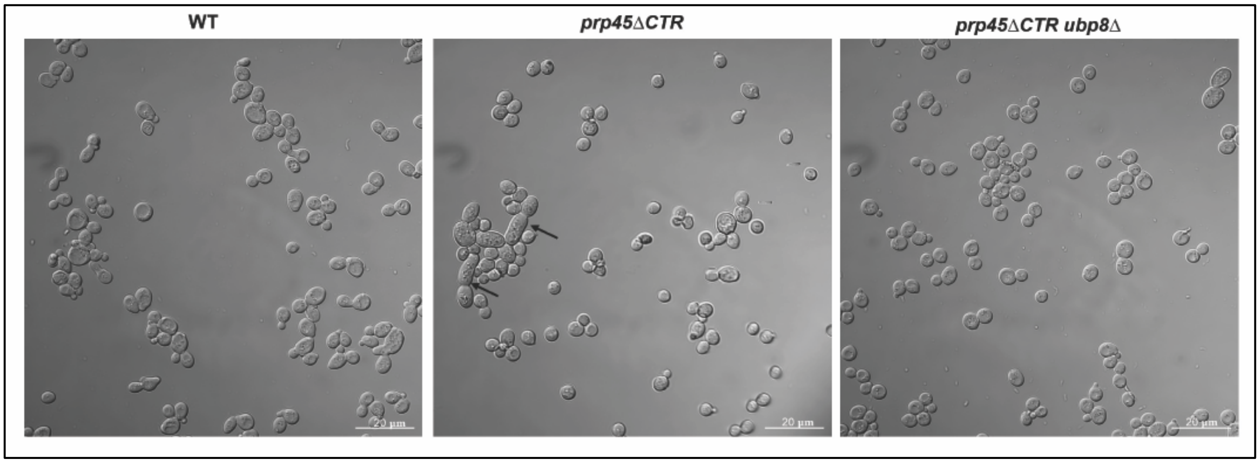


**Supplementary Figure 3. The large cell size of the *prp45ΔCTR* strain is reversed by the deletion of UBP8.** WT, *prp45ΔCTR,* and *prp45ΔCTR ubp8Δ* strains grown to $OD_{600}$ 0.6 at 37°C and Confocal imaging (DIC) was done to visualize the cell size.

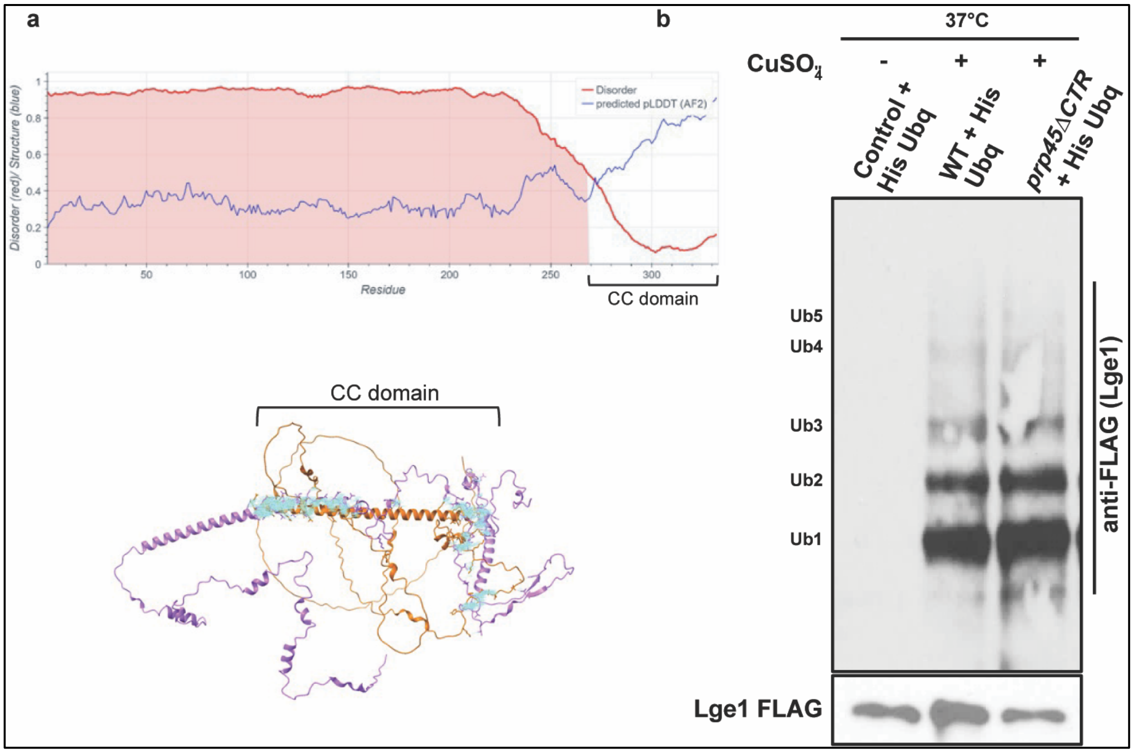


**Supplementary Figure 4. The coiled-coiled domain of Lge1 shows synthetic genetic interaction with *prp45ΔCTR.***

(a) (Upper image) The disordered and structural domains of the Lge1 protein were mapped using Metapredict. The dark shaded region indicates the disorder region, and the region from `270 to 332 is the ordered region. The sharp decrease and increase in the red and blue lines indicate that the region is more structured and a site for potential protein interaction. (Bottom) *In silico* Prp45 (purple) and Lge1 (orange) interaction analysis by AlphaFold multimer and visualized in Chimera X. The cyan color shows the interaction sites, especially at the C-terminal region of Prp45, with the Lge1. (b) Loss of C-terminal domain of Prp45 causes increased Lge1 ubiquitination. Wild type *LGE1*-FLAG and *prp45ΔCTR LGE1*-FLAG strains with histidine tagged ubiquitin expressed from the plasmid under copper inducible promoter using $CuSO_4$. Without $CuSO_4$ induction was also used as a control. Ubiquitin was pulled down with Ni-NTA agarose followed by elution with 1X SDS PAGE buffer. Ubiquitinated Lge1 was detected using anti-FLAG antibody.

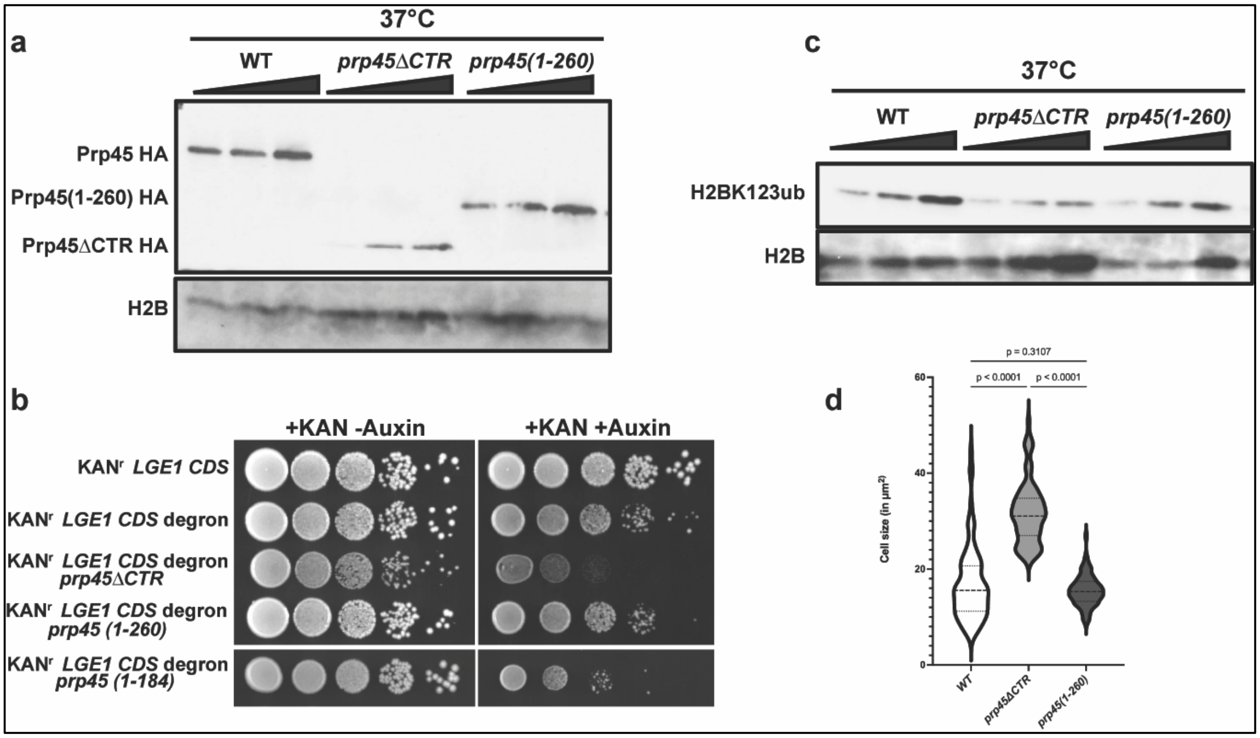


**Supplementary Figure 5. The intrinsically disordered region of Prp45 (1-260) interacts with the H2B ubiquitination machinery.**

(a) Detection of Prp45 (1-260) Whole cell lysates isolated from WT, *prp45ΔCTR,* and *prp45 (1-260)*, grown at 37°C, and analyzed by Immunoblot for the detection of Prp45, Prp45ΔCTR, and Prp45 (1-260) protein using anti-HA antibody. (b) Growth assay showing Lge1 stability, in *prp45(1-260)* and *prp45(1-184)* strains. Lge1 was N-terminally tagged with a Kanamycin resistance cassette and C-terminally tagged with a degron sequence under the native Lge1 promoter in wild type, *prp45ΔCTR* and *prp45 (1-260) and prp45(1-184)* strains. Cells were tenfold serially diluted on a YPD + kanamycin + IAA plate and incubated at 30°C. (c) Prp45 (1-260) rescues H2B monoubiquitination. Whole cell lysates isolated from WT, *prp45ΔCTR,* and *prp45 (1-260)*, grown at 37°C, and analyzed by Immunoblot for the detection of H2BK123ub and H2B using anti-H2BK123ub and H2B antibodies. (d) Prp45 (1-260) also restores the large cell morphology. WT, *prp45ΔCTR,* and *prp45 (1-260)* strains grown to $OD_{600}$ 0.6 at 37°C. Confocal imaging (DIC) was done, and cell size was measured using ImageJ. Cell size was plotted in the graph using GraphPad PRISM, and an unpaired one-way ANOVA with Tukey's post hoc analysis (n = 80) was used to calculate the *p*-values.

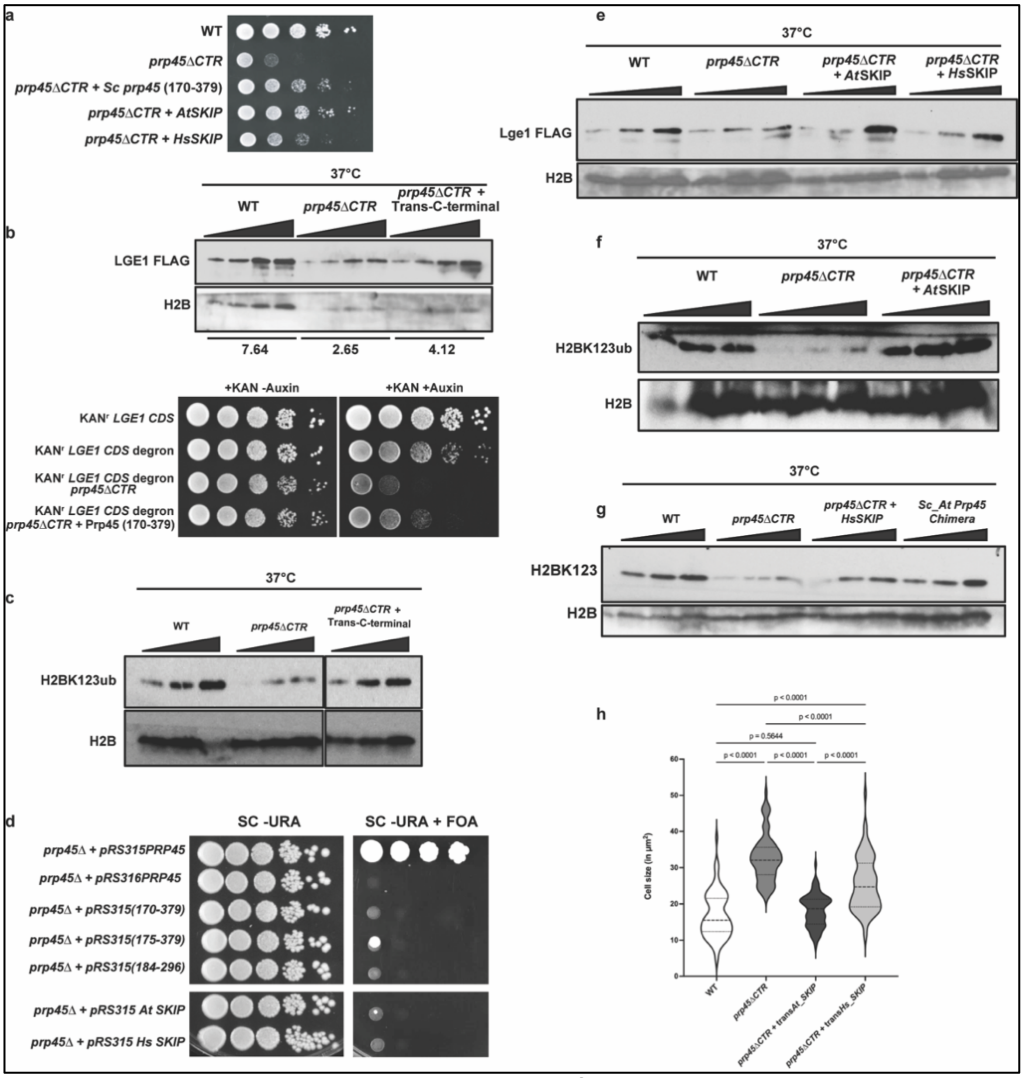


**Supplementary Figure 6**. **Prp45 homologs from metazoans support the activity of the H2B ubiquitination machinery when expressed in *S. cerevisiae***

(a) Trans expression of the C-terminal region of Prp45, full-length *At* SKIP, or *Hs* SKIP in *prp45ΔCTR* strain rescues the growth defect at 37°C. (b) Trans expression of the C-terminal domain restores the stability of the Lge1 protein. Immunoblot showing Lge1 FLAG stability from whole cell extracts of WT, *prp45ΔCTR*, and *prp45ΔCTR* + plasmid containing the C-terminal domain. H2B was used as a loading control, and densitometric quantification was done using ImageJ, as shown below.

Growth assay illustrating Lge1 stability in trans expressed C-terminal domain in *prp45ΔCTR* strains. Orthogonal degron assay was done as described in Fig.3d.

(c) Immunoblot showing H2BK123 ubiquitination from whole cell extracts of WT, *prp45ΔCTR*, *prp45ΔCTR* + C-terminal domain and H2B was used as s control.

(d) The N-terminus of Prp45 is necessary for rescue of cell growth. Different truncation mutants of the C-terminal domain of Prp45 and trans-expressed *At* or *Hs* SKIP cannot rescue the growth defect in *prp45*Δ*. prp45*Δ + pRS315 (PRP45), *prp45*Δ*C* + pRS316 (PRP45), *prp45*Δ + pRS315 (170-379), *prp45*Δ + pRS315 (175-379)*, prp45*Δ + pRS315 (184-296), *prp45*Δ + pRS315 (*At* SKIP), *prp45*Δ + pRS315 (*Hs* SKIP) cells were grown in SC-URA, tenfold serially diluted on SC FOA -URA, and incubated at 30°C for five days. (e) Trans expression of *At* SKIP and *Hs* SKIP restore Lge1 stability. Immunoblot showing the lge1's stability rescued in *prp45*Δ*CTR* strains expressing *At* or *Hs* SKIP. (f) & (g) *At* and *Hs* SKIP can rescue the H2B monoubiquitination. Immunoblot showing H2BK123 ubiquitination from whole cell extracts of WT, *prp45*Δ*CTR,* and *prp45*Δ*CTR* expressing *At* or *Hs* SKIP (S6F &G), and H2B was used as a control.

(h) *At* and *Hs* SKIP also restore yeast cell size. Confocal imaging (DIC) and image analysis followed by Cell size quantification and statistical analysis was done as mentioned before. Cell size was plotted in the graph.

**Table S1** List of proteins that show Prp45's C-terminal domain-specific interaction, and the interaction is lost when the C-terminal domain is truncated. Proteins were grouped based on their association with the major pathway complexes.

| Pathways | Specific Complex | Proteins |
|---|---|---|
| **RNA Processing** | **Spliceosome (Core & NTC)** | PRP8, PRP2, SLU7, CWC22, CWC25, CEF1, ISY1, SYF1, SYF2, BUD13, IST3, SNU66, CDC40, ECM2, FDO1 |
| **RNA Processing** | **mRNA Processing/Exosome** | HRP1, LRP1 |
| **Ribosome Biogenesis** | **SSU Processome / 90S** | KRI1, IMP4, ECM16, RIX1, GCD10 |
| **Chromatin & Transcription** | **Chromatin/Silencing (SIR)** | SIR2, SIR4, BRE1 |
| **Chromatin & Transcription** | **Transcription Factors/Stress** | REB1, HSF1, MBF1, TOD6, CTK1, SGV1 |
| **DNA Metabolism** | **DNA Replication/Repair** | MRC1, RFC5, RNR4 |
| **Nuclear/Other** | **Nuclear Transport/Mod.** | NUP1, UBA2 |
| **Nuclear/Other** | **Metabolism/Redox** | GRX1, IGD1 |

### Supplementary Material

| Genotypes | Source |
|---|---|
| *MATa his3Δ1 leu2Δ0 met15Δ0 ura3Δ0 (WT)* | Open Biosystems |
| *MATa PRP45-HA::kanMX6 LGE1-FLAG::natMX6 his3Δ1 leu2Δ0 met15Δ0 ura3Δ0* | This study |
| *MATa prp45ΔCTR-HA::kanMX6 his3Δ1 leu2Δ0 met15Δ0 ura3Δ0ura3Δ0* | This study |
| *MATa prp45ΔCTR-HA::kanMX6 LGE1-FLAG::natMX6 his3Δ1 leu2Δ0 met15Δ0 ura3Δ0* | This study |
| *MATa PRP45-HA::kanMX6 his3Δ1 leu2Δ0 met15Δ0 ura3Δ0* | This study |
| *MATa PRP45-HA::kanMX6 lge1ΔCC-FLAG::natMX6 his3Δ1 leu2Δ0 met15Δ0 ura3Δ0* | This study |
| *MATa prp45(1–174)-HA::kanMX6 his3Δ1 leu2Δ0 met15Δ0 ura3Δ0* | This study |
| *MATa prp45(1–184)-HA::kanMX6 his3Δ1 leu2Δ0 met15Δ0 ura3Δ0* | This study |
| *MATa prp45(1–260)-HA::kanMX6 his3Δ1 leu2Δ0 met15Δ0 ura3Δ0* | This study |
| *MATa prp45(1–296)-HA::kanMX6 his3Δ1 leu2Δ0 met15Δ0 ura3Δ0* | This study |
| *MATa BRE1-GFP::kanMX6 his3Δ1 leu2Δ0 met15Δ0 ura3Δ0* | This study |
| *MATa BRE1-GFP::kanMX6 prp45ΔCTR-FLAG::natMX6 his3Δ1 leu2Δ0 met15Δ0 ura3Δ0* | This study |
| *MATa ubp8Δ::natMX6 his3Δ1 leu2Δ0 met15Δ0 ura3Δ0* | Open Biosystems |
| *MATa ubp8Δ::natMX6 prp45ΔCTR-HA::kanMX6 his3Δ1 leu2Δ0 met15Δ0 ura3Δ0* | This study |
| *MATa lge1Δ::natMX6 prp45ΔCTR-HA::kanMX6 his3Δ1 leu2Δ0 met15Δ0 ura3Δ0* | This study |
| *MATa lge1ΔCC-FLAG::natMX6 PRP45-HA::kanMX6 his3Δ1 leu2Δ0 met15Δ0 ura3Δ0* | **This study** |

| | |
|---|---|
| *MATa lge1ΔCC-FLAG::natMX6 prp45ΔCTR-HA::kanMX6 his3Δ1 leu2Δ0 met15Δ0 ura3Δ* | This study |
| *MATa lge1Δ::natMX6 his3Δ1 leu2Δ0 met15Δ0 ura3Δ0e1Δ::natMX6 his3Δ1 leu2Δ0 met15Δ0 ura3Δ0* | Open Biosystems |
| *MATa bre1Δ::natMX6 his3Δ1 leu2Δ0 met15Δ0 ura3Δ00 ura3Δ0* | Open Biosystems |
| *MATa bre1Δ::natMX6 prp45ΔCTR-HA::kanMX6 his3Δ1 leu2Δ0 met15Δ0 ura3Δ0* | This study |
| *MATa rad6Δ::natMX6 his3Δ1 leu2Δ0 met15Δ0 ura3Δ0Δ0 ura3Δ0* | Open Biosystems |
| *MATa rad6Δ::natMX6 prp45ΔCTR-HA::kanMX6 his3Δ1 leu2Δ0 met15Δ0 ura3Δ0* | This study |
| *MATa PRP45-AID-HA::hphMX6 LGE1-FLAG::natMX6 his3Δ1::P(ADH1)-AtAFB2-HIS3 leu2Δ0 met15Δ0 ura3Δ0* | This study |
| *MATa sgf73Δ::natMX6 prp45ΔCTR-HA::kanMX6 his3Δ1 leu2Δ0 met15Δ0 ura3Δ0* | This study |
| *MATa sgf73Δ::natMX6 his3Δ1 leu2Δ0 met15Δ0 ura3Δ0* | This study |
| *MATa slx5Δ::HIS3MX6 his3Δ1 leu2Δ0 met15Δ0 ura3Δ0* | This study |
| *MATa slx5Δ::HIS3MX6 prp45ΔCTR-HA::kanMX6 his3Δ1 leu2Δ0 met15Δ0 ura3Δ0* | This study |
| *MATa P(LGE1)-kanR-LGE1-AID-HA::hphMX6 his3Δ1::P(ADH1)-AtAFB2-HIS3 leu2Δ0 met15Δ0 ura3Δ0* | This study |
| *MATa P(LGE1)-kanR-LGE1-AID-HA::hphMX6 prp45ΔCTR-FLAG::natMX6 his3Δ1::P(ADH1)-AtAFB2-HIS3 leu2Δ0 met15Δ0 ura3Δ0* | This study |
| *MATa P(LGE1)-kanR-LGE1-AID-HA::hphMX6 prp45ΔCTR-FLAG::natMX6 slx5Δ::URA3 his3Δ1::P(ADH1)-AtAFB2-HIS3 leu2Δ0 met15Δ0 ura3Δ0* | This study |

| Primer name | Sequence (5' to 3') | Purpose |
|---|---|---|

| | | |
|---|---|---|
| SCR1 F | AAGGGATAGTTCTCTATTCCGCACC | RT-PCR |
| SCR1 R | CACAATGTGCGAGTAAATCCTGATG | RT-PCR |
| Prp45 (Stitching forward primer promoter-300bp ) | ACG CGA GGA TCC TCG ACC GGG CAT AGA ATA GC | **Cloning & expressio n** |
| Stitching reverse primer promoter and ATG of Prp45 | GGA TTT TTC CAG TTT GAC ATA TTT GGT TGA GGT | **Cloning & expressio n** |
| Stitching forward primer for C-terminal end of Prp45 | ACC TCA ACC AAA TAT GTC AAA CTG GAA AAA TCC | **Cloning & expressio n** |
| Prp45 C-terminal stitching with FLAG sequence reverse primer | CGT CGT CCT TGT AGT C GGC GCC ATA GTT ATC CG | **Cloning & expressio n** |
| Prp45 C-terminal stitching with FLAG sequence Forward primer | CGG ATA ACT ATG GCG CC GAC TAC AAG GAC GAC G | **Cloning & expressio n** |

| | | |
|---|---|---|
| FLAG sequence + ADH1 terminator reverse primer | TTG GCC GTC GAC CCG GTA GAG GTG TGG TCA AT | **Cloning & expression** |
| Prp45 (1-169) F | AGA AGC TGA TCC AAA TGA GTG GAA GAT ACC TGC AGC TGT G CGGATCCCCGGGTTAATTAA | Deletion & tagging |
| Prp45 (1-169) R | CCA AGG CCA CGG TAT AAC CAT TTG GAT TTT TCC AGT TTG A GAATTCGAGCTCGTTTAAAC | Deletion & tagging |
| Prp45 (1-260) F | AGC CAG ATA CCA CAA CGG GAC TCC GCA GAC GGG AGC AAT A CGGATCCCCGGGTTAATTAA | Deletion & tagging |
| Prp45 (1-260) R | TTA GTC TGG CCA CTG TGC TCG TTT GCT TTT TGG GCT TAA C GAATTCGAGCTCGTTTAAAC | Deletion & tagging |
| UBP8 deletion F | GTA ATA GCA AAG GGA TTA TAT TAG TGG ACC TGA TTA ATT C CG GAT CCC CGG GTT AAT TAA | deletion |
| UBP8 deletion R | TAT AGA AAA AAG AAC GGA AGC AAG CGT TAA CTG TGA TAG G GAATTCGAGCTCGTTTAAAC | deletion |
| BRE1 deletion F | GCA TTT GAT CAC GTG ATT ATA ATC ACA TCC AAC GAA AC CG GAT CCC CGG GTT AAT TAA | deletion |
| BRE1 deletion R | TAG ACG TCA CTC TTT GAT TCA GAA GTT AGG CTA GGA TG GAATTCGAGCTCGTTTAAAC | deletion |
| SGF73 deletion F | AAA ATC ATT ACG AAC AGA AGA AAG TGG AGC TGA AGA GCG T CG GAT CCC CGG GTT AAT TAA | deletion |
| SGF73 deletion R | TGA AGA AAT ATA ATG TCT ACT TAA ATA AAT AAT GTA TAC C GAATTCGAGCTCGTTTAAAC | deletion |
| RAD6 deletion F | ATA TAC AAA AAG GCA CCT CCG GGT AGC CGG AGT AGA AAG C CG GAT CCC CGG GTT AAT TAA | deletion |

| | | |
|---|---|---|
| RAD6 deletion R | GAA TTC ATA ATA TCG GCT CGG CAT TCA TCA TTA AGA TTC T GAATTCGAGCTCGTTTAAAC | deletion |
| LGE1 C-terminal tagging F | ACT GAC CCA AGA AAA GCT GGA CTC ATT GTT ATT AAT GCA G CG GAT CCC CGG GTT AAT TAA | tagging |
| LGE1 C-terminal tagging R | TGC GTT TAC GTA GTT TAT CTA TTT ATA GGT ACG GTA TAC C GAATTCGAGCTCGTTTAAAC | tagging |
| PRP45 degron F | TAA AGCTGAATCC GATGATAAAT CGGATAACTA TGGCGCC CG TAC GCT GCA GGT CGA C | tagging |
| PRP45 degron R | AGCACAAGAATGCTTTGTTTTCCTAGTGCTCATCCTGGG CATC GAT GAA TTC GAG CTC G | tagging |
| PRP45 C-terminal tagging F | TAA AGCTGAATCC GATGATAAAT CGGATAACTA TGGCGCC CGGATCCCCGGGTTAATTAA | tagging |
| PRP45 C-terminal tagging R | AGCACAAGAATGCTTTGTTTTCCTAGTGCTCATCCTGGG C GAATTCGAGCTCGTTTAAAC | tagging |
| Lge1 degron F | ACT GAC CCA AGA AAA GCT GGA CTC ATT GTT ATT AAT GCA G CGT ACG CTG CAG GTC GAC | tagging |
| Lge1 degron R | TGC GTT TAC GTA GTT TAT CTA TTT ATA GGT ACG GTA TAC C ATC GAT GAA TTC GAG CTC G | tagging |
| HTZ1 C-terminal Turbo ID tagging F | AGC ATT ATT ATT GAA AGT GGA AAA AAA GGG AAG TAA GAA A CG GAT CCC CGG GTT AAT TAA | tagging |
| HTZ1 C-terminal Turbo ID tagging F | AGG GAG AAT TAC GGG AAA TGG GAA AGA AAA ACT ATT CTT C GAATTCGAGCTCGTTTAAAC | tagging |
| PGK1 C-terminal Turbo ID tagging F | TAA GGA ATT GCC AGG TGT TGC TTT CTT ATC CGA AAA GAA A CG GAT CCC CGG GTT AAT TAA | tagging |
| PGK1 C-terminal Turbo ID tagging R | AAA GAA AAA AAT TGA TCT ATC GAT TTC AAT TCA ATT CAA T GAATTCGAGCTCGTTTAAAC | tagging |

| | | |
|---|---|---|
| SLX5 deletion F | TGCACTAGGA GCTTCAGTCT AGGAGTGAAC TTTTGAAAAG CG GAT CCC CGG GTT AAT TAA | deletion |
| SLX5 deletion R | ATT GCA CCT CTG ATC TTT TAA TCT TAA TTA AAT AAG CGT G GAATTCGAGCTCGTTTAAAC | deletion |
| Prp45 C-terminal domain swapping with Arabidopsis F1 | AGA AGC TGA TCC AAA TGA GTG GAA GAT ACC TGC AGC TGT GTC CAA TTG GAA GAA TCC GAA AGG | **Domain swapping** |
| Prp45 C-terminal domain swapping with Arabidopsis R1 | TTA ATT AAC CCG GGG ATC CGA CGC CGG TCA CTG CGT TCG A | **Domain swapping** |
| Prp45 C-terminal domain swapping with Arabidopsis F2 | TCG AAC GCA GTG ACC GGC GTC GGA TCC CCG GGT TAA TTA A | **Domain swapping** |
| Prp45 C-terminal domain swapping with Arabidopsis R2 | CCA AGG CCA CGG TAT AAC CAT TTG GAT TTT TCC AGT TTG A GAATTCGAGCTCGTTTAAAC | **Domain swapping** |
| Prp45 C-terminal domain swapping | AGA AGC TGA TCC AAA TGA GTG GAA GAT ACC TGC AGC TGT G TC TAA CTG GAA AAA TGC AAA GG | **Domain swapping** |

| | | |
|---|---|---|
| with Human F1 | | |
| Prp45 C-terminal domain swapping with Human R1 | TTA ATT AAC CCG GGG ATC CGT TCC TTC CTC CTC TTC TTG CCT | **Domain swapping** |
| Prp45 C-terminal domain swapping with Human F2 | AGG CAA GAA GAG GAG GAA GGA ACG GAT CCC CGG GTT AAT TAA | **Domain swapping** |
| Prp45 C-terminal domain swapping with Human R2 | CCA AGG CCA CGG TAT AAC CAT TTG GAT TTT TCC AGT TTG A GAATTCGAGCTCGTTTAAAC | **Domain swapping** |
| PRP45 (1-174) C-terminal tagging F | TGA GTG GAA GAT ACC TGC AGC TGT GTC AAA CTG GAA AAA TCG GAT CCC CGG GTT AAT TAA | **Deletion & tagging** |
| PRP45 (1-174) C-terminal tagging R | TAC CTA CAC GTC TTT CCA AGG CCA CGG TAT AAC CAT TTG G GAATTCGAGCTCGTTTAAAC | **Deletion & tagging** |
| PRP45 (1-184) C-terminal tagging F | A AAC TGG AAA AAT CCA AAT GGT TAT ACC GTG GCC TTG GAA CGGATCCCCGGGTTAATTAA | **Deletion & tagging** |
| PRP45 (1-184) C- | TGG TAT TAT TTT CGT TGT CAA GAG CTT TAC CTA CAC GTC T GAATTCGAGCTCGTTTAAAC | **Deletion & tagging** |

| | | |
|---|---|---|
| terminal tagging R | | |
| PRP45 (1-296) C-terminal tagging F | G TAT CCG AAA AGA TAA TTC TGG GCG CAG CAA AGC GTT CA CGGATCCCCGGGTTAATTAA | **Deletion & tagging** |
| PRP45 (1-296) C-terminal tagging R | T TGT GAA AAA TCT TGA ATC GTA CTG CAG ATC CGG TTG TTC GAATTCGAGCTCGTTTAAAC | **Deletion & tagging** |
| Kanamycin LGE1 fusion cassette F | TTA CGA GGC AGA AGA GGT GAC CAA AAA ACC GCA GTA AAG TAT GGG TAA GGA AAA GAC TCA | **Cassette fusion** |
| Kanamycin LGE1 fusion cassette R | ATG AGT ATC TGC TAT AAT TAT TTC CCG TGT ATC CGC TCA TGA AAA ACT CAT CGA GCA TCA | **Cassette fusion** |
| At SKIP reverse stitching with Promoter | CGC AGG AAG ATC ATT AAG AGA CTT CAT ATT TGG TTG AGG T | **Cloning & expression** |
| At SKIP forward stitching with promoter | ACC TCA ACC AAA T ATG AAG TCT CTT AAT GAT CTT CCT GCG | **Cloning & expression** |
| At SKIP end stitching with HA/FLAG ADH1 terminator reverse | TTA ATT AAC CCG GGG ATC CGA CGC CGG TCA CTG CGT TCG A | **Cloning & expression** |

| | | |
|---|---|---|
| At SKIP stitching with HA/FLAG ADH1 terminator forward | TCG AAC GCA GTG ACC GGC GTC GGA TCC CCG GGT TAA TTA A | **Cloning & expression** |
| Hs SKIP reverse stitching with promoter | GGT AAA AAG CTG GTG AGC GCC ATA TTT GGT TGA GGT | **Cloning & expression** |
| Hs SKIP forward stitching with promoter | ACC TCA ACC AAA T ATG GCG CTC ACC AGC TTT TTA CC | **Cloning & expression** |
| Hs SKIP end human reverse stitching | CCT TTG CAT TTT TCC AGT TAG ACA TAT TTG GTT GAG GT | **Cloning & expression** |
| Hs SKIP end human forward stitching | ACC TCA ACC AAA TAT GTC TAA CTG GAA AAA TGC AAA GG | **Cloning & expression** |
| Hs SKIP end stitching with HA/FLAG ADH1 terminator reverse | TTA ATT AAC CCG GGG ATC CGT TCC TTC CTC CTC TTC TTG CCT | **Cloning & expression** |

| | | |
|---|---|---|
| Hs SKIP end stitching with HA/FLAG ADH1 terminator forward | AGG CAA GAA GAG GAG GAA GGA ACG GAT CCC CGG GTT AAT TAA | **Cloning & expressio n** |
